# Vertebrate extracellular matrix protein hemicentin-1 interacts physically and genetically with basement membrane protein nidogen-2

**DOI:** 10.1101/2021.11.24.469833

**Authors:** Jin-Li Zhang, Stefania Richetti, Thomas Ramezani, Daniela Welcker, Steffen Lütke, Hans-Martin Pogoda, Julia Hatzold, Frank Zaucke, Douglas R. Keene, Wilhelm Bloch, Gerhard Sengle, Matthias Hammerschmidt

## Abstract

Hemicentins are large proteins of the extracellular matrix that belong to the fibulin family and play pivotal roles during development and homeostasis of a variety of invertebrate and vertebrate tissues. However, bona fide interaction partners of hemicentins have not been described as yet. Here, applying surface plasmon resonance spectroscopy and co-immunoprecipitation, we identify the basement membrane protein nidogen-2 (NID2) as a binding partner of mouse and zebrafish hemicentin-1 (HMCN1), in line with the formerly described essential role of mouse HMCN1 in basement membrane integrity. We show that HMCN1 binds to the same protein domain of NID2 (G2) as formerly shown for laminins, but with an approximately ten-fold lower affinity and in a competitive manner. Furthermore, immunofluorescence and immunogold labelling revealed that HMCN1/Hmcn1 is localized close to basement membranes and in partial overlap with NID2/Nid2a in different tissues of mouse and zebrafish. Genetic knockout and antisense-mediated knockdown studies in zebrafish further show that loss of Nid2a leads to similar defects in fin fold morphogenesis as the loss of Laminin-α5 (Lama5) or Hmcn1. Finally, combined partial loss-of-function studies indicated that *nid2a* genetically interacts with both *hmcn1* and *lama5*. Together, these findings suggest that despite their mutually exclusive physical binding, hemicentins, nidogens, and laminins tightly cooperate and support each other during formation, maintenance, and function of basement membranes to confer tissue linkage.

## Introduction

Basement membranes (BMs) are specialized thin and dense sheets of self-assembled extracellular matrix (ECM) that are present in all animal species to link or separate cells and/or tissues. Core components of BMs are laminins, collagen IV, nidogens and the heparan sulfate proteoglycans perlecan and agrin. Together, they form complex and insoluble protein meshworks around or underneath cells or tissues (1–5). In addition, BMs contain so-called matricellular proteins (6, 7) which, in contrast to the core components, are not required for BM assembly or architecture per se, but provide additional, often tissue-specific BM properties (3). Among them are different members of the fibulin family of ECM proteins (8, 9), including their largest members, the hemicentins (10–13).

The work presented here particularly focusses on interactions between mouse and zebrafish laminins, nidogens and hemicentins. Laminins are heterotrimeric proteins consisting of an α-, β- and γ-subunit joined together through a long coiled-coil domain. Five α-, four β- and three γ-chains have been identified in mammals; however, of the possible combinations, only 16 have been shown to exist (14). Most laminins contain the γ1-subunit and loss of the corresponding *Lamc1* gene in the mouse leads to failed BM assembly and early embryonic lethality (15). Polymerization among laminin trimers is mediated via their N-terminal LN domains, while binding to cell surface receptors such as integrins α3β1, α6β1, α7β1 and α6β4 occurs via the C-terminal laminin-type globular (LG) domains of the α-subunit, and binding to nidogens and agrin via internal domains within the γ1 chain (16). In contrast, binding of laminins with perlecan and collagen IV is indirect, mediated by nidogen adaptors (3,17–19).

Nidogens are glycoproteins of 150 – 200 kDa encoded by a single gene in invertebrates, but two genes, *Nid1* and *Nid2*, in mammals (20, 21), and four genes, *nid1a*, *nid1b*, *nid2a* and *nid2b*, in zebrafish (22–24). All Nidogens consist of three globular domains, G1 to G3, with G1 and G2 connected by a flexible link and G2 to G3 by a rod-like segment containing a series of EGF-like repeats (see Figure 1A). The G2 mediates physical binding of nidogens to collagen IV and perlecan, the G3 domain binding to laminins (20,25–28). In addition, mammalian NID1 was shown to bind the matricellular protein fibulin-1 in a Ca^2+^-dependent manner, mediated via the C-terminal EGF-FC unit of fibulin-1, which is also present in hemicentins (8) and which consists of an array of Ca^2+^-binding EGF modules and a globular fibulin-type carboxy-terminal (FC) module (29, 30).

**Figure 1.**
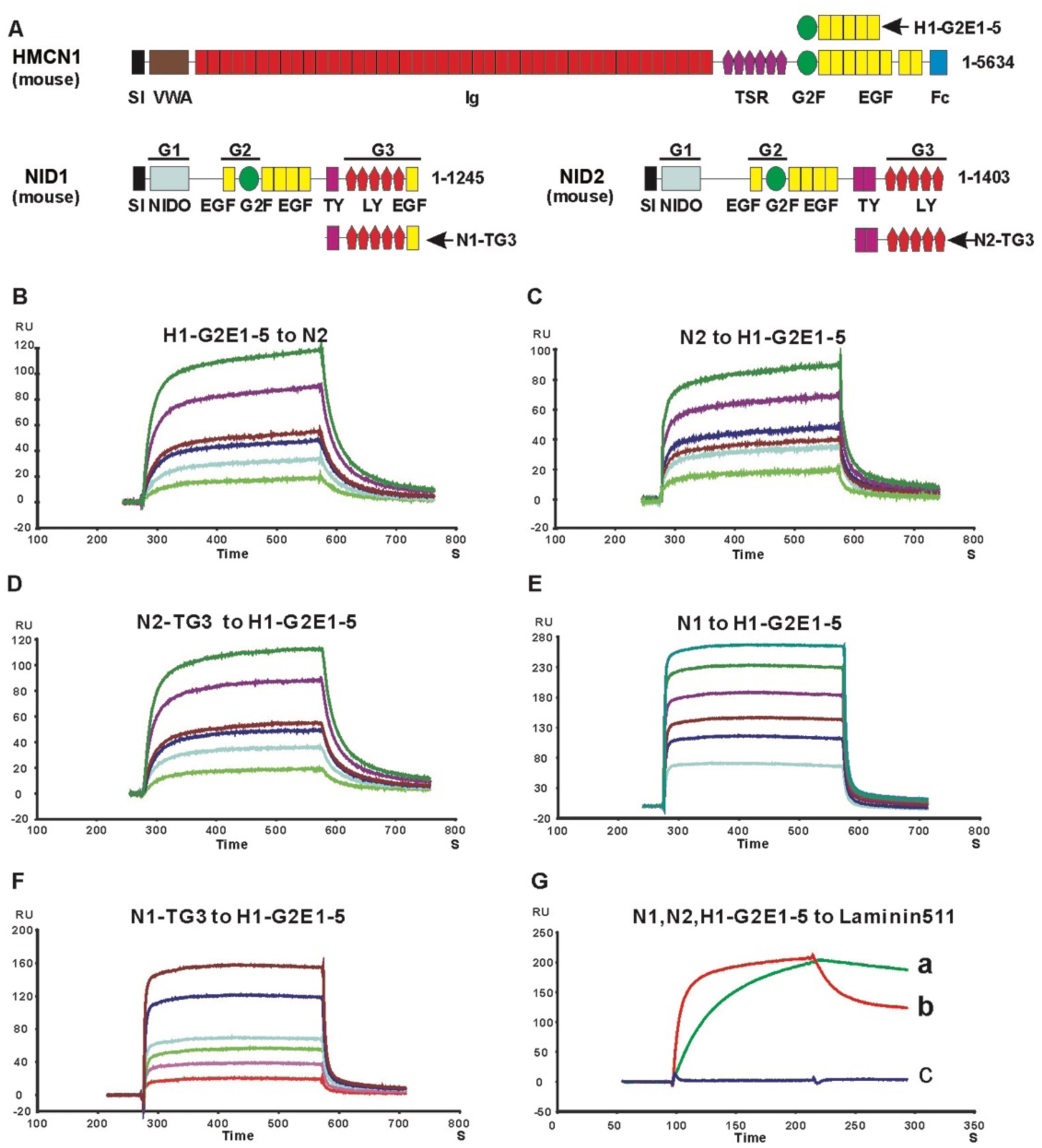
Surface plasmon resonance analysis of NID2 - HMCN1 binding. A. Schematic diagram of mouse HMCN1 and NID2 proteins and the protein fragments used for binding studies. Indicated modules are signal sequence (SI), von Willebrand type A (VWA), immunoglobulin (Ig), thrombospondin (TSR), nidogen G2F (G2F), epidermal growth factor (EGF), fibulin type carboxy-terminal (FCs), Nidogen-like (NIDO), thyroglobulin type-1(TY); and low-density lipoprotein receptor (LY) modules. The globular domains of Nidogens are indicated as G1, G2 and G3. **B-F:** Sensograms showing the binding of 10 (bottom curve), 20, 30, 40, 80, 120 (top curve) nM HMCN1 H1-G2E1-5 fragment to immobilized full-length NID2 (N2) (B), and the following proteins binding to immobilized HMCN1 H1-G2E1-5 fragment: 3 (bottom), 6, 9, 12, 24, 36 (top) nM NID2 (C); 10 (bottom), 20, 30, 40, 80, 120 (top) nM NID2-TG3 domain (D); 100 (bottom), 200, 300, 400, 600, 800 (top) nM NID1 (N1) (E); 100 (bottom), 200, 300, 400, 800, 1200 (top) nM NID1-TG3 (F). See Table 1 for K_D_ values calculated for sensograms in (B-F) **G:** NID1 (red) and NID2 (green), but not the HMCN1 fragment H1-G2E1-5 (blue), bind to immobilized laminin-511. Kinetic values are: NID1 to laminin 511, k_off_ = 2.1 x10^-3^ s^-1^, k_on_ = 8.6 x 10^+5^ M^-1^s^-1^, K_D_ = k_off_/k_on_ = 2.4 nM; NID2 to laminin 511, k_off_ = 8.1 x10^-4^ s^-1^, k_on_ = 1.6 x 10^+5^ M^-1^s^-1^, K_D_ = k_off_/k_on_ = 5 nM. Abbreviation: RU, resonance unit.

In mouse, concomitant loss of *Nid1* and *Nid2* (31, 32), or specific ablation of the nidogen binding site in the laminin-γ1 chain (33), lead to wide-spread defects attributed to regional BM ruptures, such as renal agenesis and impaired lung and heart development. In contrast, *Nid1* and *Nid2* single mutants show very selected or no overt abnormalities (34–36), pointing to largely redundant roles of the two mouse paralogs during BM assembly and maintenance. However, even in the nematode *Caenorhabditis elegans* (37–39) and in the fruitfly *Drosophila melanogaster* (40), genetic loss of their single *nidogen* gene only leads to very minor defects in BM assembly, but more prominent tissue-specific defects in BM-associated functions, such as neuromuscular junctioning and longitudinal nerve positioning in *C. elegans*. Altogether this suggests that compared to invertebrates, vertebrate nidogens have acquired additional, more global roles for BM assembly (40), acting as crucial catalysts and molecular adaptors during BM *de novo* formation and remodeling (41–43).

Hemicentins are similarly ancient and very large members (>600 kDa) of the fibulin family. In invertebrates like the *C. elegans* (in which *hemicentin* was were initially discovered, also referred to as *him-4*) (10,12,44) and planarian flatworms (45, 46), they are encoded by a single gene. In vertebrates like mammals (47–49) and zebrafish (50, 51), two paralogs exist, *Hmcn1/hmcn1*, also named *fibulin-6*, and *Hmcn2/hmcn2*. Hemicentins share a number of structural motifs (see Figure 1A): an amino-terminal von Willebrand type A (VWA) domain, followed by a long (>40) stretch of tandem immunoglobulin (Ig) domains, a G2F domain combined with multiple tandem epidermal growth factor (EGF) domains and a fibulin carboxy-terminal module (FC). Of note, the G2F-EGF modules combine to form a functional unit that apart from hemicentins is only found in nidogens, while the carboxy-terminal EGF-FC modules form a functional unit that is only found in hemicentins and in the other members of the fibulin family (fibulin1-8) (8). For *C. elegans* HMCN, the EGF-FC module has been shown to be involved in protein polymerization and track formation (52), while initial functional investigations point to roles of HMCN in transient cell contacts that are required for cell migration and BM invasion (10, 12). More recent studies identified HMCN as part of an adhesion complex that connects juxtaposed double BMs to each other, thereby conferring tissue linkage (5,53,54). Studies in planarians have further demonstrated that HMCN made by muscle cells is required for BM integrity, preventing mislocalization of neoblast stem cells and their descendants outside their normal compartments and pointing to essential roles of HMCN for proper BM function in tissue separation (45, 46). Similar roles have been suggested for mouse HMCN2 on satellite cells in skeletal muscle stem cell niches (55), while our own recent data reveal essential functions of mouse HMCN1 for proper BM integrity in dermal-epidermal junctions of the skin and in myotendinous junctions of the musculoskeletal system (49).

Previous zebrafish data from our laboratory suggest that hemicentins synergize with other ECM proteins to fulfill their essential functions during tissue linkage. We identified zebrafish *hmcn1* mutants, which show fin blistering phenotypes similar to those of mutants in *fras1* and *frem2* (50), encoding BM-associated ECM proteins mutated in human Fraser syndrome (56), and similar to those of mutants in *fibrillin2* (*fbn2*), encoding components of microfibrils mutated in human Congenital Contractural Arachnodactyly (57). In the zebrafish fin fold, such microfibrils might correspond to the so-called cross fibers that connect the two juxtaposed skin BMs and that even penetrate the BMs and contact the basal surface of epidermal cells (58), in line with the aforementioned matricellular and BM-linking roles of HMCN in invertebrates (5,53,54). Genetic interaction studies with specific antisense morpholino oligonucleotides (MOs) injected at low doses further revealed a synergistic enhancement of fin defects upon combined partial loss of *hmcn1, fras1, frem2 or fbn2* compared to partial loss of either of them alone. These data strongly suggest that *hmcn1*, *fras1*, *frem2* and *fbn2* functionally interact during zebrafish fin development *in vivo* to allow proper anchorage of the dermis and thereby tissue linkage (50). In addition, Hmcn1 seems to be required for proper interaction between the skin BM and the overlying epidermis, with mutants displaying compromised epidermal integrity, although less severe than in *lama5* mutants lacking the laminin α5 chain or in *itga3* mutants lacking its α3 integrin receptor subunit (50, 59).

However, the molecular basis of such functional interactions remained unclear. Indeed, as to now, no physical binding partners of hemicentin proteins have been reported in any species. Here we studied binding between mouse HMCN1 and different BM components, including laminins, collagen IV, NID1, NID2 and perlecan. Of those, only NID2 displayed strong binding, involving the G2F-EGF domain of HMCN1 (in contrast to the EGF-FC domain in the case of fibulin-1 binding to NID1; see above) and the G3 domain of NID2. Binding of HMCN1 to NID2 occurs in competition with laminin-γ1, but with kinetics that – together with the dynamics of HMCN1 and NID2 protein distributions *in vivo* – suggest that the HMCN1-NID2 interaction might be temporally restricted, with HMCN1 catalyzing early steps of BM assembly. Consistent with this notion, loss-of-function studies in zebrafish reveal that *nid2a* mutants display similar defects during fin fold morphogenesis as *hmcn1* and *lama5* mutants, affecting both the BM-dermal and the BM-epidermal junction. In addition, synergistic enhancement studies with combined partial loss-of-function scenarios reveal a genetic / functional interaction between Hmcn1 and Nid2a and between Nid2a and Lama5 *in vivo*, but not between Hmcn1 and Lama5 *in vivo*, which is in line with their biochemical binding properties *in vitro*.

## Results

### Mouse and zebrafish HMCN1 protein binds to nidogens

HMCN1 is a large extracellular matrix protein of over 500 kDa consisting of more than 50 domains (Figure 1A). To identify protein binding partners of HMCN1, we chose a candidate testing approach with different recombinant BM proteins. Generation of recombinant full-length HMCN1 protein failed, most likely due to its enormous size. Therefore, we instead expressed different fragments of mouse HMCN1 that altogether span 90% of the whole protein. HMCN1 fragments consisting of the VWA domain; Ig domains 8-13, 14-20, 21-26, 27-36, 33-38 or 39-44; TSR domains 1-6; the G2F domain, or a combination of subsequent TSR6-G2F-EGF1-2, G2F-EGF1-5 or EGF6-8-Fc domains (Figure 1A) were isolated and purified to >95% purity. Employing surface plasmon resonance (SPR) technology, binding of such HMCN1 fragments to collagen IV, collagen V, perlecan-Ig2-9 and - Ig10-15, laminin-111, laminin-511, nidogen-1 (NID1) and nidogen-2 (NID2) was tested. Of those, only the two nidogens were found to bind HMCN1 fragments G2F-EGF1-5, TSR6-G2F-EGF1-2 and G2F (Figure 1 and Table 1). However, binding of the HMCN1 fragment G2F-EGF1-5 to NID2 (*K_D_* of 20 nM) was approximately 30 times stronger than binding to NID1 (*K_D_* = 560 nM). Experiments with additional HMCN1 fragments for further binding domain dissections were therefore concentrated on HMCN1 and NID2. These additional SPR studies revealed that HMCN1 fragment G2F-EGF1-2 binds NID2 with an affinity similar to that of TSR6-G2F-EGF1-2 (Table 1), indicating that the TSR6 domain does not participate in binding. Furthermore, affinities of the G2F domain (*K_D_* > 3 µM) and the EGF1-5 domains (*K_D_* = 500 nM) per to Nid2 were much lower than that of the G2F-EGF1-5 fusion (*K_D_* = 20 nM). Finally, among the 5 consecutive EGF modules of the EGF1-5 domain, EGF2 and EGF5 appear most important, with EGF2 increasing the binding affinity to NID2 15 times (G2F-EGF1-2 compared to G2F-EGF1) and EGF5 increasing it 10 times (G2F-EGF1-5 compared to G2F-EGF1-4), respectively (Table 1). Together, this suggests that EGF2 and 5 of HMCN1 may form a composite interface with G2F to generate high affinity to NID2. Furthermore, binding of G2F-EGF1-5 to NID1 and NID2 strictly depends on the presence of Ca^2+^ (Table 1), suggesting that at least some of the EGF modules are Ca^2+^-binding (60), consistent with former findings for the binding of Nidogens to the C-terminal EGF-FC unit of fibulin-1 (29, 30).

**Table 1.** Dissociation constants (*K_D_* values) between mouse HMCN1 fragments and full-length or G3 domains of mouse NID1 and NID2

| HMCN1 fragments | NID2 | NID2-TG3 | NID1 | NID1-TG3 |
| --- | --- | --- | --- | --- |
| TSR6-G2F-EGF1-2 | 160 nM | 176 nM | -- | -- |
| G2F | > 3 μM | > 3 μM | -- | -- |
| G2F-EGF1 | > 3 μM | > 3 μM | -- | -- |
| G2F-EGF1-2 | 200 nM | 185 nM | -- | -- |
| G2F-EGF1-3 | 210 nM | 170 nM | -- | -- |
| G2F-EGF1-4 | 193 nM | 180 nM | -- | -- |
| G2F-EGF1-5 | 20 nM | 30 nM | 560 nM | 585 nM |
| G2F-EGF1-5 (no $\text{Ca}^{2+}$) | ND | -- | ND | -- |
| EGF1-5 | 500 nM | 620 nM | -- | -- |

--: Not determined; ND: not detectable

Nidogens themselves are multi-domain proteins consisting of NIDO, EGF, G2F, TY and LY modules that form three globular domains called G1, G2 and G3 (Figure 1A), each of which has been shown to bind to different BM proteins (20). To further dissect the binding domain of nidogens for HMCN1, we expressed G1, G1-G2, G2 and TY-G3 (TG3) domains of mouse NID1 and NID2 and tested their binding to HMCN1 fragments. As shown in Figure 1 and Table 1, the TG3 domain of NID1 and NID2 binds to HMCN1 fragments with affinities similar to those of full-length NID1 and NID2. In contrast, no binding was found between the HMCN1 fragments and the other domains of nidogens, indicating that the binding site of nidogens for HMCN1 is restricted to its TG3 domain.

But what are the reasons for the 30-fold difference in the affinities of TG3 domains of NID1 and NID2 to HMCN1? NID1 has one TY domain and a globular propeller domain followed by EGF6, which appears to be partially integrated within the propeller structure (61), while NID2 has two TY domains and lacks EGF6 (see Figure 1A). To dissect the impact of the rod-like TY domains versus the propeller G3 domain of NID1 and NID2 on HMCN1 binding, and since the “naked” propeller domains of nidogens are unstable ((28); and own data, not shown), we performed domain swapping experiments, combining the single TY domain of NID1 with the G3 domain of Nid2 (N1T-N2G3) and the two TY domains of NID2 with the G3 domain of NID1 (N2T-N1G3). In SPR studies (Supplementary Figure 1), both hybrid fragments displayed intermediate affinities to the HMCN1 G2F-EGF1-5 fragment, 4-6-fold lower than that of homotypic NID2 fragment (N2TG3), but 3-5-fold higher than that of the homotypic NID1 fragment (N1TG3). Also, N1T-N2G3 (K_D_=110 nM) displayed a 1.5-fold higher affinity than N2TN1G3 (K_D_=170 nM). Together, these data indicate that both the rod-like TY domains and the globular G3 propeller domain of the nidogens contribute to their binding - and to the different affinities of NID1 and NID2 - to HMCN1.

The interaction of the HMCN1 G2F-EGF1-5 fragment with full-length NID2 and its TG3 domain was further confirmed by co-precipitation experiments. To this end, Strep-tagged NID2 or NID2-TG3 was co-transfected with His_6_-tagged HMCN1-G2F-EGF1-5 (H1-G2E1-5) into EBNA-293 cells. The Step-tagged proteins could be co-precipitated with His_6_-tagged H1-G2E1-5 by Ni2+-NTA resin (Figure 2A,B). In a reverse experimental setup, His_6_-tagged H1-G2E1-5 could be co-precipitated with Strep-tagged NID2 or NID2-TG3 by Strep-tactin–Sepharose (Figure 2C,D). Identical results were obtained with recombinant mouse HMCN1 and NID2 fragments that had been purified from the supernatants of singly transfected EBNA-293 cells before the co-precipitation (Supplementary Figure S2), indicating that binding was direct and not mediated by other secreted EBNA-293 cell-derived proteins.

**Figure 2.**
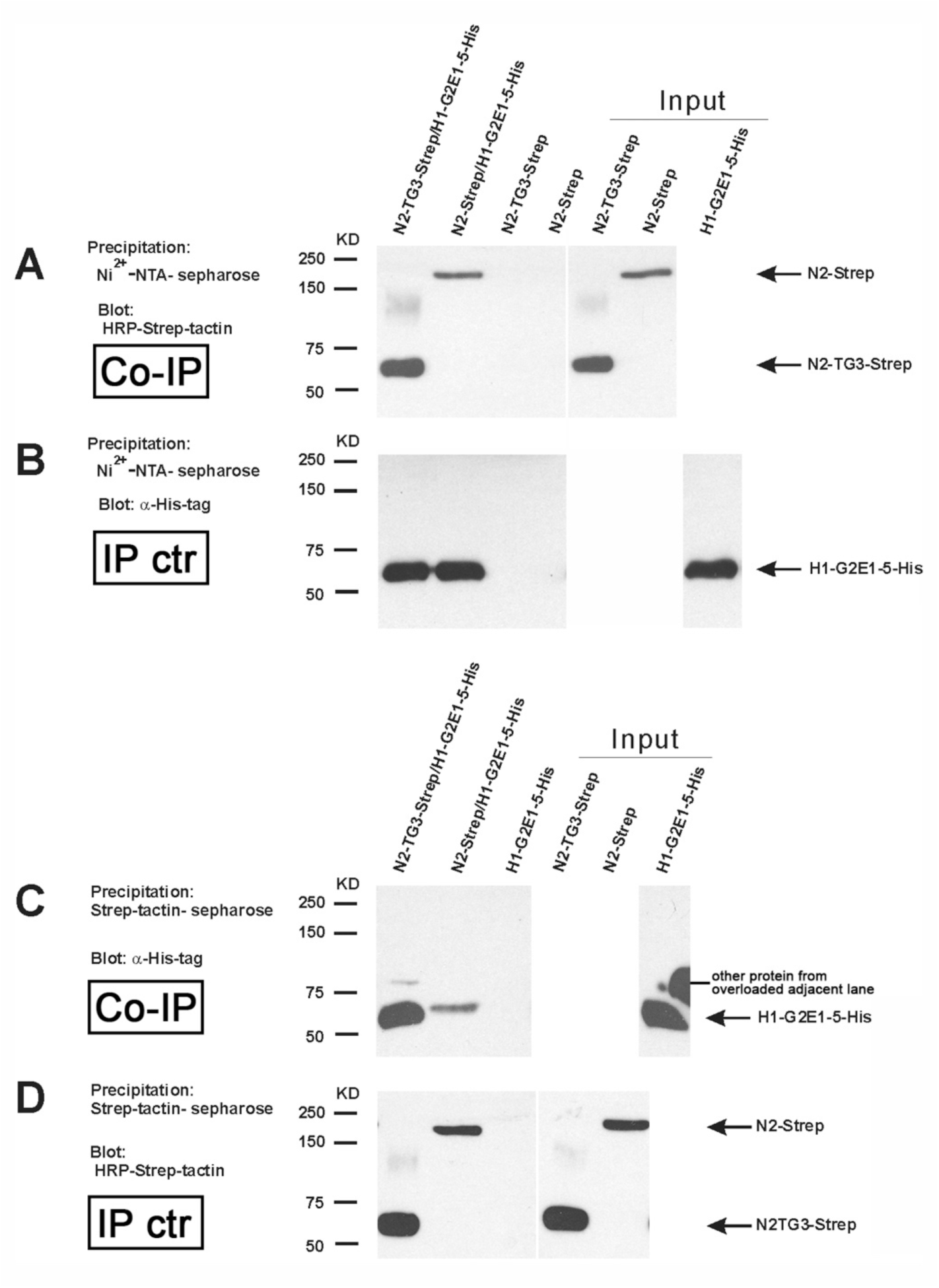
Analysis of NID2 - HMCN1 binding by co-precipitation. (A,B) Conditioned medium of cultured HEK293-EBNA cells transfected with NID2-Strep/HMCN1-G2E1-5-His_6_ or NID2-TG3-Strep/HMCN1-G2E1-5-His_6_ was precipitated by Ni^2+^-NTA sepharose and subjected to immunoblotting to detect co-precipitated Strep-tagged proteins with HRP-Strep-tactin (A) or, as control of the precipitation, His_6_-tagged proteins with anti-His_6_-tag antibodies (B) (lanes 1 and 2). As negative control, conditioned medium from cultured HEK293-EBNA cells transfected with NID2-strep or NID2-TG3-strep alone was precipitated by Ni^2+^-NTA sepharose and detected by immunoblotting (lanes 3 and 4). (C,D) For the reverse co-precipitation, conditioned medium of cultured HEK293-EBNA cells transfected with NID2-Strep/ HMCN1-G2E1-5-His or N2-TG3-Strep/ HMCN1-G2E1-5-His was precipitated by Strep-tactin-sepharose and subjected to immunoblotting to detect co-precipitated His_6_-tagged proteins with anti-His_6_-tag antibodies (C) or, as control of the precipitation, Strep-tagged proteins with HRP-Strep-tactin (D) (lanes 1 and 2). As negative control, conditioned medium from cultured HEK293-EBNA cells transfected with HMCN1-G2E1-5-His_6_ alone was precipitated Strep-tactin-sepharose and detected by immunoblotting (lane 3). Lanes 5-7 of (A-B) and 4-6 of (C,D) are direct loadings of the indicated proteins (50% of input). Abbreviations: Co-IP, co-immunoprecipitation; IP ctr, control immunoprecipitation.

To look into the evolutionary conservation of this binding, and to better relate the identified biochemical interactions with the functional zebrafish investigations described below, we also performed SPR and co-precipitation studies with the corresponding purified recombinant zebrafish protein fragments (Supplementary Figures S3 and S4). Both assays revealed direct binding between the 2TY-G3 fragment of zebrafish Nid2a and the G2-E1-5 fragment of zebrafish Hmcn1, with - according to the SRP data – a K_D_ value of 30 nM, thus an affinity as that of the corresponding mouse fragments (30 nM; see above, Figure 1D and Table 1).

### HMCN1 competes with laminin for binding to the G3 domain of nidogen-2

It is known that via their γ-chain short arms, laminins can bind to the G3 domain of NID1 and NID2 (25). Actually, while according to former reports, the affinity of NID1 to Laminin is much higher than that of NID2 (approximately 1000-fold for human nidogens (62) and 20-fold for mouse nidogens (63)), our own data obtained for the TG3 fragments of the mouse proteins point to more similar affinities (NID1: K_D_ = 2.4 nM; NID2: K_D_ = 5 nM; Figure 1G). As the HMCN1 binding site of NID2 is also located in TG3, we wondered how the three proteins interact in concert with each other, and therefore performed competitive SPR analyses. Strikingly, mouse H1-G2E1-5 could inhibit the binding of NID2 to laminin-γ1 short arm (LAMγ1) immobilized on the chip in a concentration-dependent fashion (Figure 3A). In a reverse experimental set-up, LAMγ1 short arm inhibited the binding of Nid2 to immobilized H1-G2E1-5, whereas a N836D LAMγ1 short arm amino acid exchange mutant deficient in nidogen binding (64) could not inhibit NID2-HMCN1 binding (Figure 3B). Of note, the affinity between the LAMγ1 short arm and NID2 (Figure 3A; K_D_ = k_off_ / k_on_ = 6 nM) was approximately 3.5-fold higher than that between the HMCN1 fragment H1G2E1-5 and NID2 (Figures 1C and 3B; K_D_ = 20 nM). This is due to an approximately 12-fold lower dissociation rate between LAMγ1 and NID2 (k_off_ = 7.3 x 10^-4^ s^-1^) compared to HMCN1 and NID2 (k_off_ = 83.7×10^-4^ s^-1^), which over-compensates the approximately 3.5-fold lower association rate (k_on_ = 1.2 x 10^5^ M^-1^s^-1^ for LAMγ1 and NID2 versus 4.2 x 10^5^ M^-1^s^-1^ for HMCN1 and NID2). Together, this indicates that laminin competes with HMCN1 for NID2 binding, adding HMCN1 as an additional player in the nidogen-laminin-integrin network that can bind NID2 more quickly, but less persistently than laminins. Of note, however, given the rather similar affinities of NID1 and NID2 to laminin, this competition of HMCN1 / laminin for NID2 binding should be much less pronounced at sites with relatively higher amounts of NID1 protein, such as the skin and muscle of adult mice (63), since the excessive NID1 out-competes nearly all NID2 attachment to laminins, leaving it free to interact with HMCN.

**Figure 3.**
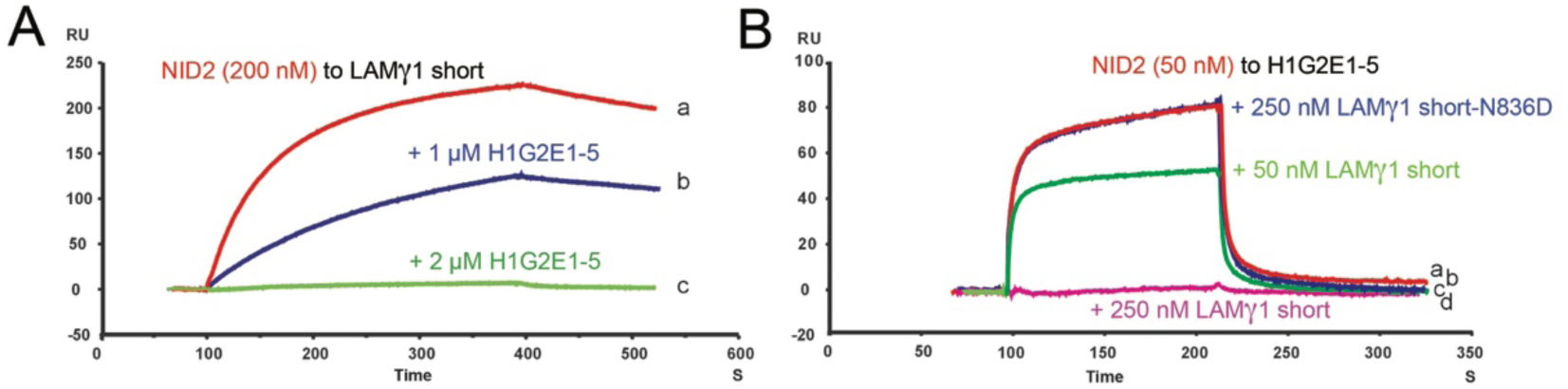
Surface plasmon resonance analysis of competition between HMCN1 and LAMγ1 for NID2 binding. LAMγ1 short arm (A) or HMCN1 fragment HMCN1-G2E1-5 (B) were immobilized on the chip surface, and the chip was perfused with 200 nM NID2 (A, a; K_D_ = k_off_ / k_on_ = 6 nM), 200 nM NID2 plus 1 µM HMCN1-G2E1-5 (A, b), 200 nM NID2 plus 2 µM HMCN1-G2E1-5 (A, c), 50 nM NID2 (B, a, red), 50 nM NID2 plus 250 nM LAMγ1 mutant deficient in nidogen binding mutant (N836D; (81, 82) (B, b, blue), 50 nM NID2 plus 50 nM LAMγ1 short arm (B, c, green), or 50 nM NID2 plus 250 nM LAMγ1 short arm (B,d, magenta), respectively.

### Mouse and zebrafish HMCN1/Hmcn1 proteins are localized closely adjacent to basement membranes, partially overlapping with NID2/Nid2a

As a first step to study whether HMCN1 and NID2 also interact *in vivo*, we performed immunofluorescence analyses to determine the spatial distribution of the two proteins in mouse and zebrafish. Former immunofluorescence studies with specific antibodies have revealed mouse HMCN1 protein in mesenchymal tissues such as the dermis of the skin and tendons of the musculoskeletal system, while its paralog HMCN2 is localized in the epithelial counterparts such as epidermis of the skin and the endomysium of skeletal muscle (49). Double immunofluorescent labeling of skin sections with HMCN1 and NID2 antibodies (Figure 4A-C) and anti-HMCN1 immunogold labeling (Figure 4D,E) further showed that in particular along follicles of epidermal appendages such as whisker follicles (Figure 4A-C) and hair follicles (Figure 4D,E) dermal HMCN1 protein accumulates directly underneath (Figure 4A,E) or even within the skin BM, partially overlapping with NID2 (Figure 4C). Similarly, endomysial HMCN2 is localized closely adjacent to NID2 and LAMγ1 in skeletal muscle (Figure 4F-H).

**Figure 4.**
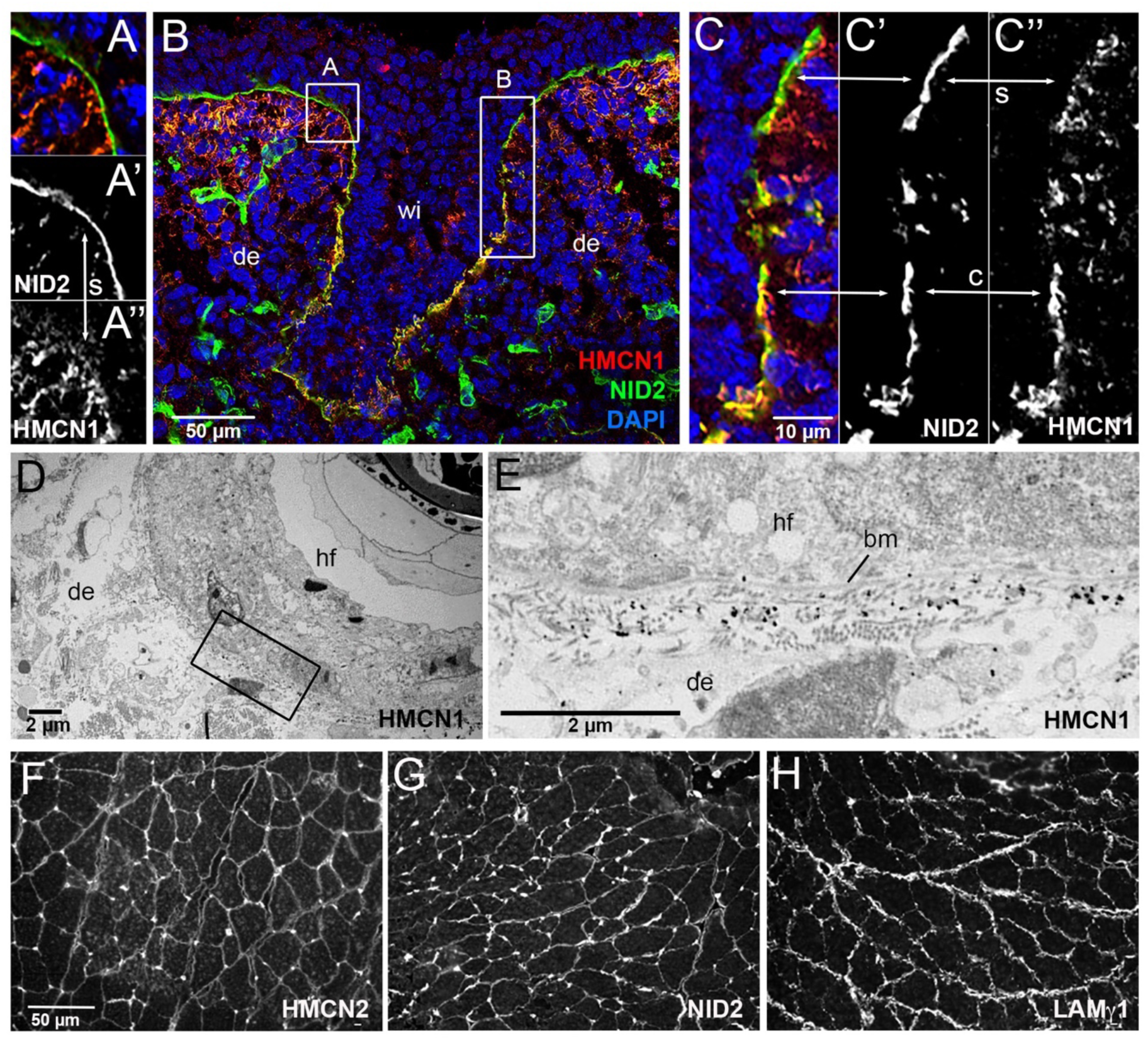
HMCN1 and NID2 proteins display partially overlapping localization at basement membranes of mouse embryos. (A-C) Immunofluorescence analysis of skin at the level of the whisker follicle of embryo at E14, using antibodies directed against HMCN1 (red) and NID2 (green); the section was counterstained with DAPI to label nuclei (blue). Merged (A,B,C) as well as single NID2 (A’,C’) and single HMCN1 (A’’,C’’) channels are shown. (A,C) show magnifications of regions boxed in B. In apical regions of the whisker follicle, dermal HMCN1 protein has accumulated underneath the BM, largely, but not completely, separated from NID2 located within the BM (A,A’,A’’,C,C’,C’’; indicated by arrows labelled with s). In contrast, in deeper regions of the whisker follicle, HMCN1 is largely located within the BM, co-localizing with NID2 (insets C,C’’,C’’; indicated by arrows labelled with c). (D) Overview of ultrastructural immunolocalization of HMCN1 protein on skin sections of P14 mouse. (E) Higher magnification reveals clusters of HMCN1 signals underneath the BM. (F-H) Consecutive transverse sections of musculus gastrocnemius from wild-type adult mouse immunofluorescently labelled with antibodies against HMCN2 (F), NID2 (G) and LAMγ1 (H). HMCN2, NID2 and LAMγ1 display similar distributions along endomysial BMs of muscle fibers. Although not resolved in these immunofluorescence labelings, we assume that HMCN2 is localized between the two adjacent BMs of neighboring muscle fibers, partially overlapping with BM nidogens, similarly to how it is shown below for Hmcn1 in embryonic zebrafish myotendinous junctions (Figure 5J-L). Abbreviations: bm, basement membrane; c, co-localized; de, dermis; hf, hair follicle; s, separated; wi, whisker.

In zebrafish, *hmcn1* has been shown to be expressed in distal epidermal cells of the embryonic fin folds, where it is required for proper dermal-epidermal junction formation in developing median fin folds (50). In addition, whole mount in situ hybridization at 30 hours post fertilization (hpf) revealed a thus far undescribed expression in tenocyte precursor cells (Figure 5E), which at these embryonic stages are localized lateral to the somitic muscle, extending cellular protrusions into the somitic myosepta to contribute to myotendinous junctions (65, 66). Consistently, specific antibodies raised within the frame of this work (see Experimental Procedures) revealed Hmcn1 protein both in the dermal space between the two epidermal sheets of the median fin folds (Figure 5G,H), closely associated with the skin BM (Figure 5M,N), as well as in the myosepta (Figure 5G-L).

**Figure 5.**
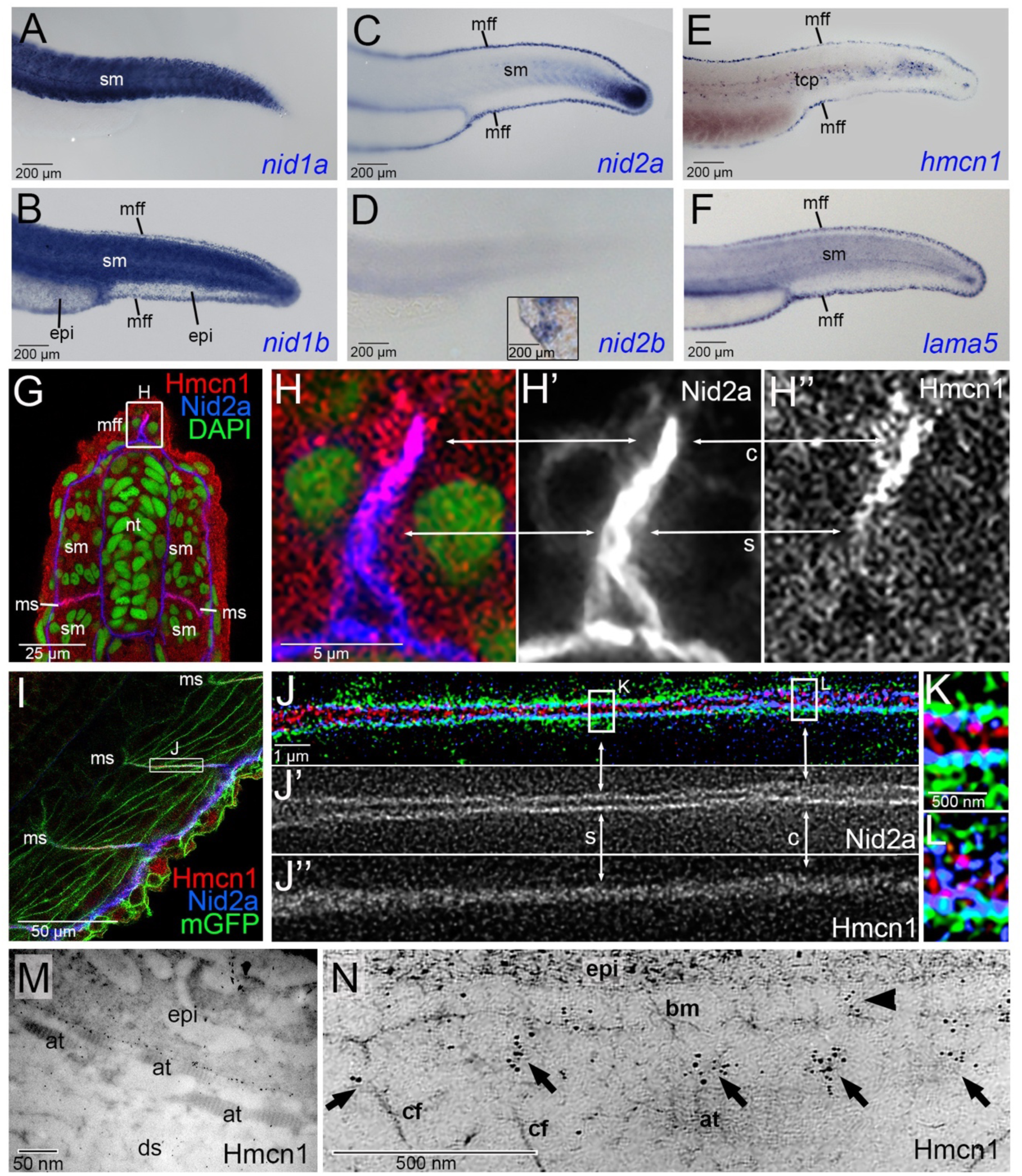
Hmcn1 and Nid2a proteins display partially overlapping localization at basement membranes of zebrafish embryos. (A-F) Lateral views of tails of zebrafish embryos at 30 hpf, stained via whole mount in situ hybridization for *nid1a*, *nid1b*, *nid2a*, *nid2b*, *hmcn1* or *lama5* transcripts. *hmcn1* and *nid2a* as well *as lama5* are co-expressed in epidermal cells of the median fin fold and in somitic muscle cells. Note the expression of *hmcn1* transcript in tenocyte precursor cells (tcp, E). Inset in D shows labelling of hatching gland cells of same embryo, serving as a positive *nid2b* control. (G-L) Immunofluorescence analysis of developing dermal-epidermal (G-H’’) and myotendinous junctions (I-L) of 24 hpf zebrafish, using antibodies directed against Hmcn1 (red) and Nid2 (blue), and counterstained with DAPI (green) to label nuclei (G,H) or with an antibody against *Tg(Ola*.*Actb*:*Hsa*.*hras-egfp)*-encoded GFP to label cell membranes (I-K). (H-H’’) shows higher magnification of the apical region of dorsal median fin fold boxed in G, (J-J’’) shows higher magnification of myotendinous junction region boxed in (I), and (K,L) further magnifications of regions boxed in (J). (G,H,I,J,K,L) show merged channels, (H’,J’) Nid2a single channels and (H’’,J’’) Hmcn1 single channels. (H-H’’) Higher magnification of the median fin fold shows Hmcn1 localization largely co-localized with Nid2a in distal, developmentally less advanced regions (50) of the fin fold (arrows labelled with “c”). In comparison, in proximal, developmentally more advanced regions, Hmcn1 and Nid2 localization have started to segregate (arrows labelled with “s”). (J-J’’) Higher magnification of the mytendinous junction between two myosepta using STED microscopy shows Hmcn1 localization largely separated (indicated by arrows labelled with “s” in J and magnified in K) or co-localized (indicated by arrows labelled with “c” in J and magnified in L) with Nid2a protein. (M) Overview of ultrastructural immunolocalization of Hmcn1 at dermal-epidermal junction of median fin fold of 54 hpf embryo. (N) Higher magnification reveals clusters of Hmcn1 signals distributed underneath (arrows) or even within (arrowhead) the skin BM. Abbreviations: at, actinotrichia; bm, basement membrane; c, co-localized; cf, cross fibers; ds, dermal space; epi, epidermis; mGFP, cell membrane-associated GFP; mff, median fin fold; ms, myoseptum; s, separated; sm, somitic muscle; tcp, tenocyte precursors.

Of the four *nidogen* genes, *nid1a*, *nid1b*, *nid2a* and *nid2b* (ZFIN Zebrafish Information Network; ZDB-GENE-050302-58, ZDB-GENE-070802-3, ZDB-GENE-030827-2, ZDB-GENE-040724-143), whole mount in situ hybridization revealed that *nid1b* and *nid2a*, but not *nid1a* and *nid2b*, are co-expressed with *hmcn1* and the laminin-α5 gene *lama5* in epidermal cells of the median fin fold (Figure 5A-F). In addition, *nid1a*, *nid1b* and *nid2a* are expressed by somitic muscle cells (Figure 5A-D). Furthermore, co-immunofluorescence studies with specific antibodies raised against zebrafish Nid2a (see Materials and Methods) revealed partially overlapping localization of Nid2a and Hmcn1 proteins both in the tip of the median fin fold (Figure 5G,H) and in the myosepta (Figure 5I-L). In the median fin fold, co-localization was particularly evident in distal regions, where new Hmcn1 and Nid2a proteins are made, while in more proximal regions, the Hmcn1 and Nid2a domains have started to segregate (see arrows in Figure 5H). Similar regions of protein co-localization next to regions of protein segregation were revealed via STED microscopy for the myosepta. Here, regions with segregated proteins showed Nid2a located within the two juxtaposed skin BMs directly adjacent to corresponding two muscle cell membranes, but Hmcn1 between these two BMs, possibly linking them (Figure 5I,J). Together, these results indicate that at least during early stages of dermal-epidermal and myotendinous junction formation, Nid2 and Hmcn1 proteins are localized in a partially overlapping manner.

### Loss of zebrafish Nid2a, Lama5 or Hmcn1 cause similar defects during median fin fold development

To address whether hemicentins and nidogens interact functionally *in vivo*, we carried out comparative and combined loss-of-function studies. Mouse *Hmcn1* mutants only show very mild defects in dermal-epidermal and mytendinous junctions (49), while defects do not seem further enhanced upon combined loss of *Hmcn1* and *Hmcn2* (48). This largely dispensable nature of mouse hemicentins precludes comparative and combined hemicentin-nidogen loss of function studies, and we therefore turned to zebrafish. Here, loss-of-function mutations in both *hmcn1* and the laminin-α5 chain gene *lama5* have been shown to cause compromised dermal-epidermal junction formation in the embryonic median fin fold (50). Loss of Hmcn1 primarily affects the connection of the skin BM with the underlying dermis, causing cutaneous blistering. In comparison, the linkage between the skin BM and the overlying epidermis is less affected, only leading to moderately compromised cohesion among basal keratinocytes during more advanced stages of skin development (48 hpf) (50). This is in contrast to the defects caused by loss of integral BM components as in *lama5* mutants, which display compromised linkage to overlying keratinocytes, leading to severe disintegration of the epidermis (50), as well as compromised linkage to the underlying dermis, leading to strong dermal blistering, the main phenotypic trait of *hmcn1* mutants (compare panels Q, S and T of Figure 6).

**Figure 6.**
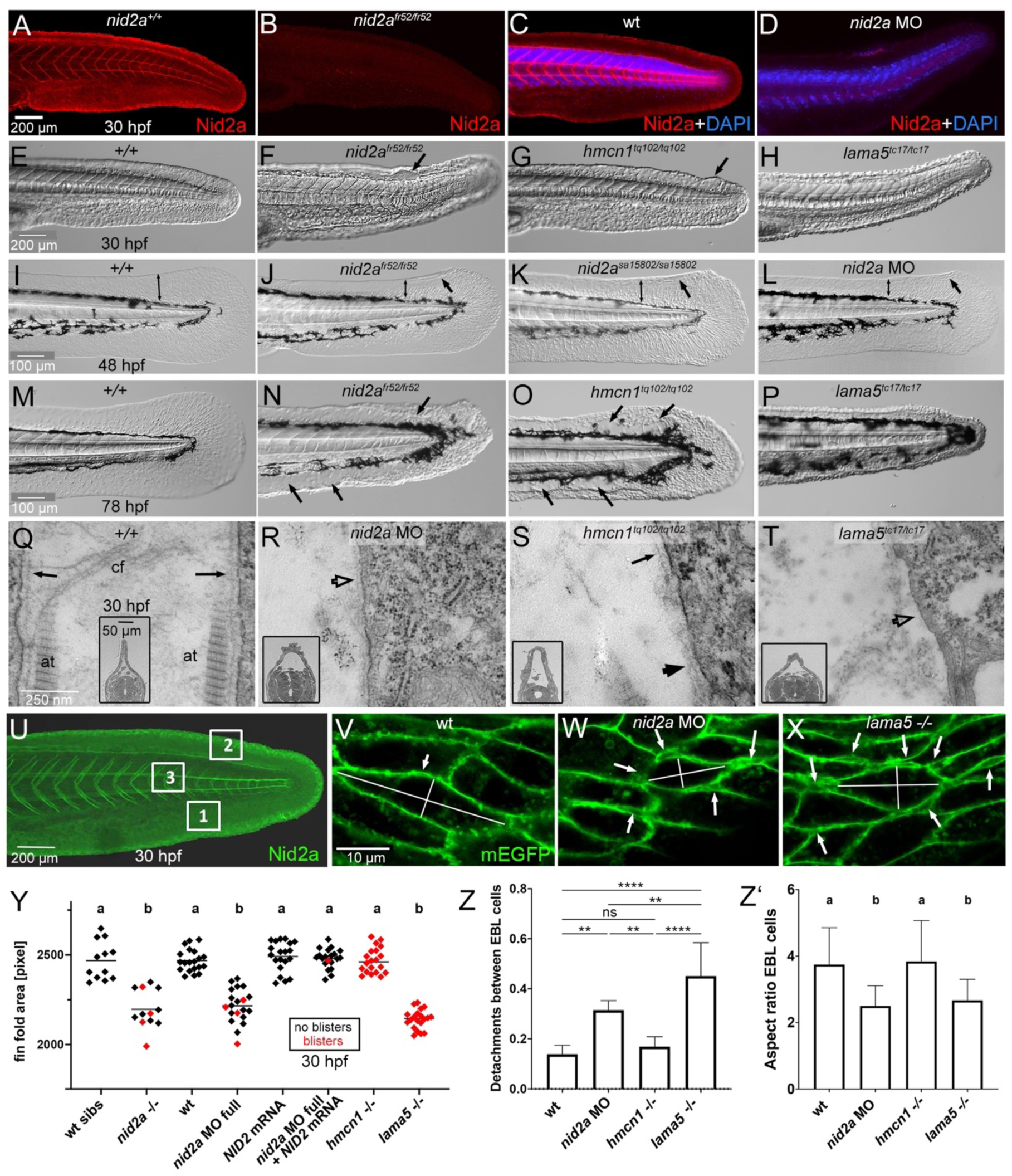
Median fin fold defects of *nid2a* morphant / mutant zebrafish embryos in comparison to *hmcn1* and *lama5* mutants. (A-D) Immunofluorescence labelling of Nid2a protein, in C,D counterstained with DAPI to label nuclei, of *nid2a^fr52^* mutant (B), full *nid2a* morphants (D) and the respective wild-type controls (A,C), 30 hpf, lateral view on tails. For full *nid2a* knockdowns, also shown in panels L,R,W,Y-Z’, 2.2 ng *nid2a-1.ATG* MO was injected per embryo (2.5 nl of 0.1 mM solution). (E-H) Lateral Nomarski images of tails / median fin folds of wild-type, *nid2a^fr52^*, *hmcn1^tq102^* or *lama5^tc17^* mutant embryos, 30 hpf. Arrows point to blisters. (I-L) Lateral Nomarski images of tails / median fin folds of wild-type, *nid2a^fr52^*, *nid2a^sa15802^* and full *nid2a* morphant, 48 hpf. Bilateral arrows demarcate extension of dorsal median fin folds, regular arrows point to blisters. (M-P) Lateral Nomarski images of tails / median fin folds of wild-type, *nid2a^fr52^*, *hmcn1^tq102^* or *lama5^tc17^* mutant embryos, 78 hpf. Arrows point to blisters. (U-X) Transmission electron micrographs of dermal-epidermal junctions in dorsal median fin fold of wild-type, *nid2a* morphant, *hmcn1^tq102^* or *lama5^tc17^* mutant embryo, 30 hpf, transverse section through tail region; inset show low magnification overviews of dorsal halves of sections, also illustrating the increased width of the dermal space between the two adjacent epidermal sheets of the dorsal median fin fold in *nid2a* morphant and *hmcn1* and *lama5* mutants. Arrows point to BM of normal morphology, empty arrow heads in R,T point to regions with indistinct BM in *nid2a* morphant and *lama5* mutant, and filled arrowhead in S points to regions with widened BM in *hmcn1* mutant, consistent with the recently reported BM defects in *Hmcn1* mutant mice (49). (U) Lateral view on tail region of wild-type embryo, 30 hpf, after immunofluorescent labeling of Nid2a (green); boxed regions were used to perform the analyses of keratinocyte detachments and cell shape changes shown in panels V-X,Z,Z’. Regions 1 and 2 are ventral and dorsal median fin fold areas, respectively, and prone to blister formation; region 3 is within body axis and not prone to blistering. (V-X) Representative regions 1 of wild-type, *nid2a* morphant and *lama5* mutant embryo, 30 hpf, double-transgenic for *Tg(Ola*.*Actb*:*Hsa*.*hras-egfp)^vu119Tg^*, labeling membranes of all cells with EGFP (green), and *Tg(krt4:tomato-CAAX)^fr48Tg^*, specifically labeling membranes of outer layer epidermal cells with Tomato to distinguish them from epidermal basal layer (EBL) cells (red channel not shown). Arrows point to detachments between basal keratinocytes of the EBL, summarized in panel Z, lines indicate the longest and shortest axes of representative cells used to determine the aspect ratios (degree of cellular elongation) summarized in panel Z’. (Y) Scatter plot of median fin fold sizes, indicating compromised BM-epidermal linkage in 30 hpf *nid2a^fr52^*, *hmcn1^tq102^* or lama5*^tc17^* mutant and in 30 hpf wild-type embryos injected with the indicated MOs and/or mRNAs. Each diamond represents one individual embryo. Red diamonds represent embryos with blisters, indicating compromised BM-dermal linkage. Note that co-injection of mouse *Nid2* mRNA rescues both the median fin fold size and the blistering defects of *nid2a* morphant embryos, proving the specificity of the *nid2a* MO. (Z,Z’) Column bar graphs quantifying numbers of detachments between basal keratinocytes of the EBL, normalized against numbers of investigated keratinocytes (Z) and the elongation degree (ratio between longest and shortest axis; Z’) of such basal keratinocytes of wild-type, *nid2a* morphant, *hmcn1* mutants and *lama5* mutant embryos, 30 hpf, scoring the three tail regions marked in panel U and images as shown in panels V-X. Average ratios and standard deviations are indicated. Significances of differences in Y-Z’ were determined via ANOVA followed by the Tukey’s post-hoc test. In U,W, groups with different superscript letters (a,b) are significantly different (p<0.0001). In V, **** and ** indicate significant differences with p<0.0001 and p<0.0070, respectively. ns indicates no significant differences. Abbreviations: at, actinotrichia; cf, cross fibers; EBL, epidermal basal layer.

Loss-of-function studies of zebrafish *nidogen* genes had thus far only been reported in the context of eye development and embryonic body size determination, but not for median fin fold development (22–24). Using CRISPR/Cas9 technology, we generated a genetic null mutant (*nid2a^fr52^*) in *nid2a*, which of the four zebrafish *nidogen* genes shows an expression pattern most similar to that of *hmcn1*, including, together with *nid1b*, a shared expression in the median fin fold (see above; Figure 5). In addition, we studied the formerly described mutant allele *nid2a^sa15802^* (23, 24) and also knocked down *nid2a* in wild-type embryos with morpholino antisense oligonucleotides (MOs) to generate *nid2a* morphants. Mutants and morphants displayed similar phentypes: anti-Nid2a immunofluorescences studies, which yielded strong Nid2a labelling in wild-type or un-injected siblings, failed to detect Nid2a protein in *nid2a^fr52^* mutant and *nid2a* morphant embryos, indicating efficient knockout and knockdown (Figure 6A-D).

Similar to *lama5* mutants, *nid2a* mutants and morphants display progressively reduced sizes of the median fin folds at 30 hpf, 48 hpf and 78 hpf (Figure 6E-P, Y), indicative of a progressive collapse of that structure as a morphological consequence of epidermal disintegration. Using transgene-encoded GFP-labelling of the cell membranes of basal keratinocyte, such compromised epidermal integrity could be directly visualized, revealing increased numbers of basal keratinocyte detachments from each other in *nid2a* morphants and *lama5* mutants already at 30 hpf (Figure 6U-X,Z). Additionally, a rounding-up of basal keratinocytes was observed in *nid2a* morphants and *lama5* mutants, yet not in *hmcn1* mutants at this early time point (Figure 6U-X,Z’). These epidermal defects most likely result from compromised BM-epidermal junction formation. In addition, *nid2a* mutants and morphants display cutaneous blistering similar to that of *hmcn1* and *lama5* mutants, most likely resulting from compromised BM-dermal junction formation (Figure 6E-P,Y). This is particularly evident in transmission electron microscopy of transverse sections (Figure 6Q-T), revealing i) a widening of the dermal space between the two epidermal sheets of the fin fold (see also Figure 7A-H), ii) the absence of dermal collagenous fibers known as actinotrichia directly underneath the cutaneous BM of *nid2a* morphants, *hmcn1* mutants and *lama5* mutants, iii) reduced BM material in *nid2a* and *lama5* mutants, and dermally disintegrated BM material in *hmcn1* mutants. Of note, this blistering / BM-dermal junction trait of *nid2a* morphants is less severe and less penetrant than in *hmcn1* and *lama5* mutants (Figure 6Y). However, it is of similar magnitude to *lama5* mutants when *nid1b*, the other *nidogen* gene expressed in median fin fold keratinocytes, is knocked down in addition to *nid2a* via injection of dedicated MOs (Supplementary Figure S5), indicating that for establishing a linkage with the underlying dermis, *nid2a* acts in partial functional redundancy with *nid1b*.

**Figure 7.**
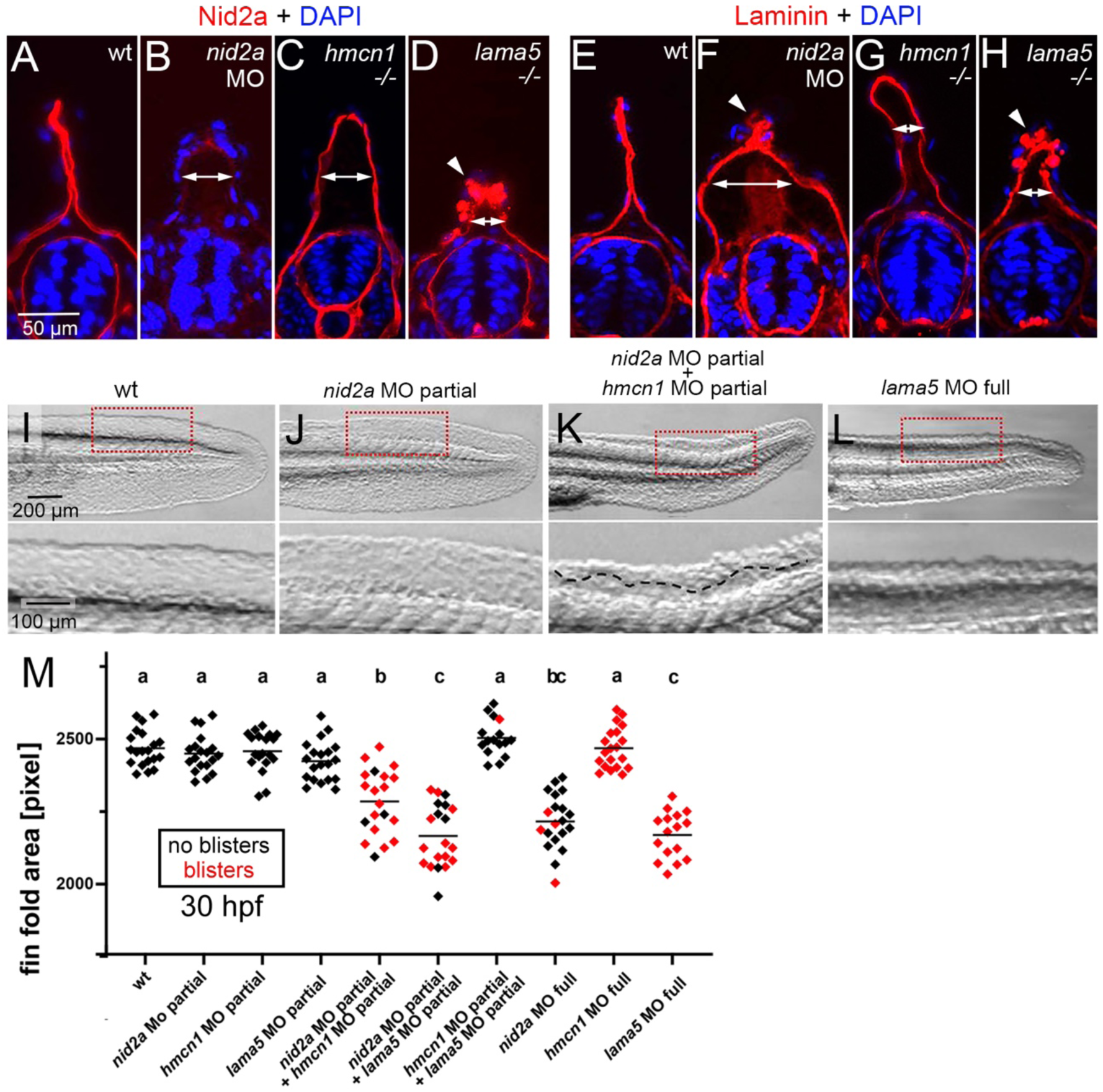
Hmcn1, Nid2a and laminin do not stabilize each other at the protein level but display genetic interaction *in vivo*. (A-H) Immunofluorescence labelling of Nid2a (A-D) and Laminin (E-H) (red) of wild-type siblings, *nid2a* morphants, *hmcn1* mutants and *lama5* mutants at 30 hpf, counterstained with DAPI (blue) to mark nuclei, transverse sections through tail region. Arrows indicate the degree of widening of the dermal space between the two cutaneous BMs of the fin fold labeled by Nid2a or Laminin, indicative of compromised BM-dermal linkage. Arrowheads point to ectopic Nid2a or Laminin within the epidermis, most likely resulting from compromised BM and epidermal epithelial integrity. (I-L) Lateral Nomarski images of tails / median fin folds of representative embryos summarized in M, 30 hpf, after injection with the indicated single or combined MOs. Lower panels show magnified views of region boxed in upper panels. Dashed line in lower panel of K demarcates region with blister. (M) Quantification of median fin fold sizes (indicative of compromised BM-epidermal linkage) in embryos at 30 hpf after injection with the indicated single or combined MOs. Each diamond represents one individual embryo. Diamonds representing embryos with blisters, indicative of compromised BM-dermal linkage, are in red. Injected amounts of MOs per embryo were: *nid2a* MO full, 2.2 ng; *nid2a* MO partial, 1.1 ng; *hmcn1* MO full, 4.3 ng; *hmcn1* MO partial, 1.1 ng; *lama5* MO full, 0.44 ng; lama5 MO partial, 0.11 ng. For *lama5* MO full and *hmcn1* MO full, compare with Figure 6Y for *lama5 -/-* and *hmcn1 -/-* null mutants. Significances were determined via ANOVA followed by the Tukey’s post-hoc test. Groups with different superscript letters (a,b,c) are significantly different (a-b and a-c, p<0.0001; b-c, p<0.0020).

Together, these data show that, despite the competitive binding of Hmcn1 versus Laminin to Nid2 *in vitro* (see above, Figure 3), all three seem to cooperate to contribute to BM-epidermal as well as to BM-dermal linkage *in vivo*.

### Hmcn1, Nid2a and Laminin proteins are not required for mutual stabilization

Members of the Fraser complex, BM-associated extracellular proteins also involved in BM-dermal linkage (50,56,67), have been shown to stabilize each other (50, 68). To investigate whether the same might be true for Hmcn1, Nid2a and Laminins, we carried out mutual immunofluorescence studies in *nid2a* morphants and *hmcn1* or *lama5* mutants, revealing altered fin fold shapes and, to some extent, altered protein distribution patterns of Nid2a or laminin. However, labeling intensities of Nid2a in *hmcn1* and *lama5* mutants, and of Laminin in *hmcn1* and *nid2a* mutants (Figure 7A-H) were unaltered, indicating that zebrafish Hmcn1 is not required for Nidogen-2 and Laminin protein stabilization, consistent with its formerly shown dispensability for Fras1 protein stability (50). Furthermore, zebrafish Nid2a is not required for Laminin stability, consistent with former immunofluorescence data for laminin-γ1 in mouse *Nid2* mutants (36). Of note, ectopic pericellular Laminin was also observed around keratinocytes in the tips of the median fin folds of *nid2a* morphant and *lama5* mutant zebrafish embryos (Figure 7F,H, arrowheads), most likely resulting from the compromised integrity of the skin BM and the epidermis observed in the TEM analyses described above (Figure 6Q-Z’).

### Zebrafish Hmcn1 genetically interacts with Nid2a during embryonic median fin morphogenesis

Injection of lower amounts of MOs allows to generate a continuous series of hypomorphic conditions. This technique is widely used in zebrafish genetics to investigate whether combined partial knockdown of two genes has synergistically enhancing effects, indicative of a functional interaction of the two gene products *in vivo* (see (51) for our former data on Hmcn1, Fras1 and Fibrillin-2). Upon injection of one quarter of the MO amount required for full knockdown and phenotypes identical to those of null mutants, *nid2a*, *hmcn1* and l*ama5* partial morphants displayed wild-type morphology (normal fin fold sizes and no blistering; Figure 7I,J,M). However, when combined, *hmcn1* / *nid2a* double partial morphants as well as *nid2a* / *lama5* double partial morphants displayed strong fin fold size reduction (resulting from compromised BM-epidermal linkage) and strong dermal blistering (resulting from compromised BM-dermal linkage), comparable to the defects of full *nid2a* and full *lama5* morphants. In contrast *hmcn1* / *lama5* double partial morphants showed wild-type fin fold morphology (Figure 7I,L,M), consistent with formerly published data (50). This synergistic enhancement of defects in the *hmcn1* / *nid2a* and *nid2a* / *lama5* combinations indicates that the corresponding partners display a strong genetic and functional interaction *in vivo*. Although a functional interaction does not necessarily require a physical interaction of the corresponding protein, it js strikingly consistent with the biochemical binding between mammalian HMCN1 and NID2 and between zebrafish Hmcn1 and Nid2a revealed here (Figures 1, 2 and Supplementary Figures 2-4), and between NID2 and laminins shown elsewhere (16). Furthermore, the absence of such a genetic interaction between Hmcn1 and Lama5 is consistent with the lack of physical binding between HMCN1 fragments and LAMγ1 in our surface plasmon resonance studies (Figure 1), suggesting that their cooperation to promote BM linkage occurs in a more indirect manner (see Discussion).

## Discussion

### HMCN1 and NID2 physically bind each other *in vitro*

With a molecular weight of over 500 kDa, HMCN1 is the largest member of the fibulin family of ECM proteins, consisting of more than 50 domains (Figure 1A). Our previous studies showed that mutations in the zebrafish *hmcn1* gene result in embryonic fin blistering (50), most likely due to compromised linkage between the two adjacent epidermal BMs of the embryonic median fin folds and comparable to the essential role of HMCN as a BM-LINKage component connecting adjacent tissues through adjoining BMs in the nematode *C. elegans* (53, 54). More recently, we have shown that loss of HMCN1 in mice results in a widening and compromised integrity of the BM in dermal-epidermal and myotendinous junctions (49), comparable to the BM defects observed upon loss of function of hemicentin in planarians (46). However, the exact alterations within the ECM network underlying such BM integrity and linkage defects in hemicentin-deficient invertebrates and vertebrates remained unclear.

The domains of HMCN1 belong to types known to be involved in the binding among ECM / BM or between ECM / BM and cell surface proteins, suggesting that HMCN1 interacts with several other proteins to fulfill its matricellular functions. Other members of the fibulin family (8, 69) have been shown to physically interact with multiple proteins, including cell surface receptors like integrins (70) and the epidermal growth factor receptor (EGFR) (71), other matricellular proteins like fibronectin (72), BM proteins like laminins and nidogens (29) and fibrillary proteins like fibrillins and tropoelastin (73, 74). In contrast, although discovered over twenty year ago (10), no physical binding partners of hemicentins had been reported in any animal species thus far.

Here, testing multiple BM proteins via SPR studies, we identified nidogens as binding partners of mouse HMCN1 (Figures 1,2; Table 1). Of note, however, NID2 binds Hmcn1 with an approximately 30x higher affinity than NID1. This is in striking contrast to former results obtained for fibulin-1 (FBLN1), which only binds NID1 (28–30), but not NID2 (62). Furthermore, we identified the G2F-EGF unit of HMCN1 (which is absent in FBLN1) to be involved in NID2 binding, whereas NID1 – FBLN1 binding is mediated via the EGF-FC unit of FBLN1 (which is also present in HMCN1). Yet, both biochemical interactions are Ca^2+^-dependent. Ca^2+^-binding of cbEGF-domains has been shown to be essential for proper folding and protein-protein interactions (75), and our SPR data also suggest that the presence of Ca^2+^ is required to guarantee structural support of cbEGF-2 and cbEGF-5 for proper binding of the G2F-EGF unit of HMCN1 to NID2.

Of note, *C. elegans* HMCN and FBLN-1 have been shown to recruit each other to BM-associated locations (11), while zebrafish Hmcn2 and Fbln1 have been shown to play redundant roles during dermal-epidermal junction (DEJ) formation (51). Together, this suggests that despite their different direct binding partners, NID1 and NID2 may display comparable co-operations with the HMCN – FBLN1 system.

Strikingly, of all annotated proteins, apart from hemicentins, a G2F-EGF unit as identified here as HMCN1’s binding domain to the G3 domain of NID2, is only present in nidogens themselves, where it mediates binding of NID1 and NID2 to collagen IV and perlecan (20,25,27,28). Theoretically, this newly identified G2F – G3 binding could enable several further protein-protein interactions. First, via their G2F-EGF units, hemicentins could also bind collagen IV and perlecan. However, both were found negative in our initial SPR screen for binding to the different mouse Hmcn1 fragments. Second, nidogens could display intermolecular or intramolecular binding between their G2 and G3 domains, thereby driving nidogen polymerization / fibril formation and/or attenuating G2-mediated binding to perlecan or collagen IV and G3-mediated binding to laminins. Along these lines, occupancy of NID2’s G3 domain by HMCN1 would prevent nidogen polymerization and promote nidogen binding to perlecan or collagen IV. However, in our SPR assays, no physical binding between G2 and G3-bearing fragments of mouse nidogens was detected (JLZ and MH, unpublished observations), making such scenarios rather unlikely.

In contrast, we have demonstrated that HMCN1 binding to NID2’s G3 domain has implications to its physical interaction with laminins (25, 26). Thus, binding of HMCN1’s G2F-EGF to NID2’s G3 domain occurs in direct competition with LAMγ1, which is present in most laminin trimers (Figure 3). Strikingly, although the overall thermodynamic affinity of NID2 to HMCN1 is approximately 3.5 times lower than that to LAMγ1, the kinetic association constant between NID2 to HMCN1 is approximately 3.5 times higher than that between NID2 and LAMγ1, but over-compensated the 12 times higher dissociation constant. These complex *in vitro* binding properties suggest that HMCN1 has a more sophisticated role in the BM niche. NID2 – HMCN1 binding may be rather temporary, and it is tempting to speculate that HMCN1 may act as a kind of chaperone during early steps of BM assembly, for instance allowing the G2-mediated interaction of NID2 to perlecan or collagen IV before its G3-mediated binding to laminins.

### HMCN1/Hmcn1 and NID2/Nid2a co-localize *in vivo*

Such a temporary function of HMCN1 – NID2 binding during early steps of BM assembly would be consistent with the dynamics of HMCN1/Hmcn1 – NID2/Nid2a localization observed in mouse and zebrafish embryos, respectively. Our former analysis with chimeric embryos have revealed that during zebrafish median fin fold development (58), BM proteins are generated by epidermal cells at the tip of the evaginating fin folds. As the fin fold grows, these cells continue to move further distally while leaving BM material behind to constitute dermal-epidermal junctions (DEJs) in more basal regions of the fin folds (50). Thus, DEJs in basal regions of the fin folds should be developmentally more advanced than in distal regions. Of note, such distal / developmentally less advanced DEJs show Hmcn1 – Nid2a co-localization, while in basal / developmentally more advanced DEJs the domains the distributions have started to segregate, with Nid2a localized within the BM and Hmcn1 directly underneath it (Figure 5). The same phenomenon was observed in DEJs along the (invaginating) whisker follicles of mouse embryos, with co-localization of HMCN1 and NID2 protein at the tips of the invaginations, but largely separated distributions at the base of the invaginations, again most likely representing developmentally less and more advanced DEJs, respectively (76, 77) (Figure 4). Together, this suggests that the relative distributions of HMCN1/Hmcn1 and NID2/Nid2a change during the time course of BM assembly, with co-localization during early phases (when it may act as a chaperone of nidogen-laminin binding, see above), but spatial segregation during later phases.

Future studies have to reveal potential additional binding partners of HMCN1 during such later stages, when HMCN1 is largely located underneath, rather than directly within the BM (Figures 4 and 5), possibly mediating proper anchorage of the BM to the underlying tissues. Binding partners with such anchoring functions seem likely, given that in zebrafish, *hmcn1*, *nid2a* and *lama5* mutants display compromised BM-dermal junction formation in the embryonic skin. Thus, the dermal actinotrichia, thin ray-like structures composed of fibrillary collagens, have left their positions directly underneath the BM (Figure 6Q-T), suggesting that Hmcn1 might also bind actinotrichial collagens, bridging the BM with the underlying collagen fibrils. Furthermore, in comparison to *nid2a* and *lama5* mutants, *hmcn1* mutants are largely characterized by dermal blistering between the two skin BMs of the fin folds, indicative of compromised BM-dermal-BM linkage, rather than a fin fold collapse, indicative of compromised BM integrity (Figure 6U-Z’). This points to binding partners of Hmcn1 mediating cross-linking of adjacent BMs, as formerly described in invertebrates (5,53,54), possibly involving thus far unidentified components of the cross fibers of embryonic zebrafish fin folds (58).

In addition, it will be interesting to look for potential hemicentin binding partners at cell surfaces mediating their “matricellular” functions (3, 78) and, at least in *C. elegans*, involving HMCN’s N-terminal VWA domain (52). Thereby, especially the unknown contribution of mammalian HMCN2 deserves further investigation. Here, we observed that HMCN2 localizes close to NID2 and LAMγ1 in the endomysium of the gastrocnemius muscle of adult mice (Figure 4F-H). This suggests that HMCN2 plays a role for BM integrity in muscle tissues, consistent with a corresponding function of HMCN2 proposed for the satellite cell niche of skeletal muscle (55).

### Hmcn1 and Nid2a functionally co-operate *in vivo*

Importantly, in addition to the physical binding *in vitro* and the temporary co-localization *in vivo*, we could also demonstrate a functional co-operation between Hmcn1 and Nid2a in zebrafish embryos *in vivo*. Thus, both are required for proper formation and function of both the BM-dermal and the BM-epidermal junction (Figure 6), and both display a tight genetic interaction during dermal-epidermal junction formation, each synergistically enhancing the effect of the other upon partial loss of function (Figure 7). Of note, similar dermal-epidermal junction defects are caused by loss of function of *lama5* (Figure 6) (50, 59), encoding the alpha chain of Laminin-511, formerly known as Laminin-10, which is also crucial for mammalian skin development (79). However, in contrast to *hmcn1* and *nid2a* and in contrast to *nid2a* and *lama5*, *hmcn1* does not genetically interact with *lama5* (Figure 7). This indicates that despite their competitive binding to Nid2, Hmcn1 also co-operates, but in a more indirect manner, with laminins to promote BM assembly and functioning, in line with the aforementioned chaperone model.

It is also noteworthy that the DEJ defects caused by loss of *hmcn1*, *nid2a* or *lama5*, although strikingly similar, are not completely identical. For the BM-dermal junction, the blistering caused by loss of *nid2a* is weaker than that caused by loss of *hmcn1* or *lama*5 (Figure 6Y). This is largely due to a partial functional redundancy between *nid2a* and *nid1b* during the establishment of the BM-dermal (but not the BM-epidermal) junction. Thus, only the combined loss of both paralogs leads to blistering defects as strong as those of *hmcn1* and *lama5* mutants (Supplementary Figure S5), consistent with the partially redundant roles of *Nid1* and *Nid2* in the mouse (31, 32). Also, at the ultrastructural level, mouse (49) and zebrafish *Hmcn1*/*hmcn1* mutants display a dermal extension of BM material not apparent upon loss of *nid2a* or *lama5* (Figure 6Q-X), pointing to an additional, possibly Nid2a and Lama5-independent or even Nid2a and Lama5-antagonistic role of Hmcn1 to spatially restrict BM formation. Similarly, for the BM-epidermal junction, *hmcn1* mutants develop epidermal integrity defects significantly later (48 hpf) (50) than *nid2a* morphants and *lama5* mutants (30 hpf; Figure 6Z,Z’), pointing to additional functions of laminins and nidogens that are at least partly independent of Hmcn1.

It is difficult to judge whether hemicentins and nidogens display similar functional co-operations in other organisms. In mouse, the phenotype of *Hmcn1* mutants (48, 49) is much weaker than that of *Nid1/2* double mutants (31, 32). This is consistent with the weaker phenotypes of mutants in other BM-associated matricellular proteins compared to integral BM components (7), but largely precludes functional cooperation studies *in vivo*. In *C. elegans*, loss of the single *nidogen* (37–39) or *hmcn* (10,12,53,54) genes lead to rather dissimilar defects, pointing to crucial functions of the two that are independent of each other. However, *C. elegans* HMCN and fibulin-1 proteins have been shown to recruit each other (11), and fibulin-1 has been shown to recruit nidogen (80) to shared assembly sites *in vivo*, suggesting that also in worms, nidogens and members of hemicentin/fibulin family can functionally interact (78) and that this interaction is highly evolutionary conserved.

## Experimental Procedures

### Recombinant expression and purification of mouse and zebrafish hemicentin-1 and nidogen proteins and protein fragments

The cDNAs encoding mouse HMCN1, NID1, NID2 and their fragments, the Laminin-γ1 short arm and the Laminin-γ1 short arm N836D mutant deficient in nidogen binding, as well as the cDNAs encoding the zebrafish Hmcn1 fragment zfH1-G2E1-5 and the zebrafish Nid2a fragment zfN2a-TG3 were cloned into a modified pCEP-Pu vector containing an N-terminal BM-40 signal peptide and a C-terminal double StrepII-tag downstream of the restriction sites (64,81,82). The recombinant plasmids were introduced into HEK293-EBNA cells (Invitrogen, Carlsbad, CA, USA) using FuGENE 6 transfection reagents (Roche, Basel, Switzerland). The cells were selected with puromycin (1 μg/ml) and recombinant proteins were purified directly from serum-containing cell culture medium. After filtration and centrifugation (30 min, 10,000× *g*), the cell culture supernatants were applied to a Strep-tactin column (IBA Lifesciences GmbH, Göttingen, Germany) and eluted with 2.5 mM desthiobiotin, 10 mM Tris–HCl, pH 8.0. The eluted proteins were then purified by Fractogel® EMD TMAE (Merck KGaA, Darmstadt, Germany) or SP-Sepharose (GE-Healthcare, Chicago, IL, USA) ion exchange chromatography. Perlecan Ig10-15 fragment was expressed and purified in HEK293-EBNA cells as described above, perlecan Ig2-9 fragment was kindly provided by Takako Sasaki, Oita University, Japan. Collagen IV, collagen V and laminin-111 were purchased from Sigma-Aldrich (St. Louis, MO, USA), laminin 511 from BioLamina AB (Sundbyberg, Sweden).

### Surface plasmon resonance analysis

The binding of mouse and zebrafish HMCN1 fragments to nidogens and other matrix proteins was recorded on a BIAcore 3000 system (GE-Healthcare). Protein ligands in 10 mM sodium acetate at pH 4 were coupled to the carboxymethylated dextran layer of CM5 sensorchips which were activated with 0.05 M N-ethyl-N’-(3-diethylaminopropyl)-carbodiimide and 0.05 M N-hydroxysuccinimide following a manufacturer’s protocol. Interaction was analyzed by recording sensograms in HBS buffer (10 mM HEPES, pH7.4, 150 mM NaCl, 2.5 mM CaCl_2_, 0.005% surfactant P20). Surfaces were regenerated by perfusion for 1.5 min with 4 M MgCl_2_. All measurements were corrected for non-specific interactions by subtracting a control sensogram recorded for flow cell 1. For dissociation constants (*K_D_*) of <50 nM, kinetic constants (*k*_on_ and *k*_off_) were evaluated. For *K_D_* values of >50 nM, dose-dependent equilibrium binding was evaluated. Per experiment, *K_D_* values or rate constants *k*_off_/*k*_on_ were determined for 6-9 different concentrations of the analytes. Mean values of *K_D_* and their standard deviation (S.D.) values were calculated from the values of at least three different experiments. S.D. values for the obtained affinities were less than 50%.

### Co-precipitation studies

The cDNAs encoding the mouse HMCN1 fragment H1-G2E1-5 or the zebrafish Hmcn1 fragment zfH1-G2E1-5 were subcloned into pCEP4 vector containing hygromycin resistance gene, an N-terminal BMP40 signal and a His_6_-tag downstream of the restriction site. For Figure 2, the mouse construct was transfected into HEK293-EBNA cells containing the plasmid pCEP-pu-mNID2-strep-tag or pCEP-pu-mNID2-TG3-strep-tag (both containing a puromycin resistance). The cells were selected with puromycin (1 μg/ml)/hygromycin (100 µg/ml) and the confluent cells were cultured for 3 days. The conditioned medium was recovered by centrifugation, and 1 ml of the conditioned medium was mixed with 40 μl of Ni^2+^-NTA sepharose beads (Invitrogen) at 4°C for 1h. The beads were then washed for three times with PBS, pH 7.4 at 4°C, centrifuged at 500 rpm for 5 min, and boiled in SDS-PAGE loading buffer for 3 min. The supernatant was separated into two parts and subjected to SDS-gel electrophoresis under reducing conditions and immunoblotting. NID2 proteins were detected by HRP-conjugated strep-tactin (IBA-GmbH), and H1-G2E1-5 was detected by mouse anti-His_6_-tag antibody (Invitrogen) and subsequently by HRP-conjugated secondary antibody (Jackson ImmunoResearch). As a negative control, conditioned medium from cultured HEK293-EBNA cells transfected with NID2-strep or NID2-TG3-strep alone was precipitated by Ni^2+^-NTA sepharose beads and detected by anti-His_6_-tag antibody in parallel.

To test whether His_6_-tagged H1-G2E1-5 could be co-precipitated by Strep-tagged NID2 or NID2-TG3, in a reverse setup the experiments were done in the same way, but the conditioned medium was precipitated by Strep-tactin-sepharose (IBA-GmbH) and the conditioned medium from cultured HEK293-EBNA cells transfected with His_6_-tagged H1-G2E1-5 alone was taken as a negative control.

To further verify the interaction between mouse H1-G2E1-5 and mouse NID2-TG3 and between zebrafish H1-G2E1-5 and zebrafish NID2a-TG3, respectively, and to exclude stabilization of the complexes by other secreted 293-cell derived protein(s), co-precipitations were performed with His-tagged H1-G2E1-5 and Strep-tagged N2-TG3 fragments purified from the supernatants of HEK293-EBNA cells that had been singly transfected with the HMCN1 or NID2 expression constructs (Supplementary Figures S2 and S4). Experiments were essentially carried out as described above, using 1 µg of each of the two purified protein fragments in 300 µl TBS buffer (10 mM Tris/HCl pH 7.5, 150 mM NaCl, 2.5 mM CaCl_2_) that was mixed with 20 µl Ni^2+^-NTA or Strep-Tactin-sepharose.

### Animals

For mouse experiments, the wild-type strains C57BL/6J or 129/Sv were used. The mutant zebrafish lines *hmcn1^tq102^* (50) and *lama5^tc17^* (50), and the transgenic lines *Tg(Ola*.*Actb*:*Hsa*.*hras-egfp)^vu119Tg^* (83) and *Tg(krt4:tomato-CAAX)^fr48Tg^* (84) were previously described. Mutant line *nid2a^sa15802^*, generated via ENU chemical mutagenesis, was purchased from the European Zebrafish Resource Center (EZRC, https://www.ezrc.kit.edu). It contains an A-G nonsense point mutation leading to premature termination of the Nid2a protein at aa 437 and nonsense-mediated decay of the mutant transcript, most likely representing an amorph or strong hypomorph (23). The mutant line *nid2a****^fr52^*** line was generated in this work, using CRISPR-Cas9 technology. The reverse strand of *nid2a*-exon2 from position bp73 to 92 was chosen as target site using the ZiFiT-web-tool (http://zifit.partners.org/ZiFiT/). Synthetic dsDNA of a dedicated gRNA scaffold (85) was obtained from IDT Inc. (Belgium): 5’- GCGTAATACGACTCACTATAGGCGATGACGGGGAAGTCGGTGTTTTAGAGCTAGAAATAGCAAGTTAAAATAAGGCTAGTCCGTTATCAACTTGAAAAAGTGGCACCGAGTCGGTGCTTTAAACGCG-3’. This *nid2a*-exon2 gRNA template was cloned via TA-cloning into the pCR^TM^II vector (Invitrogen). Sequence verified plasmid-DNA was linearized by *Dra*I digest, PCI extracted following standard protocols and subsequently transcribed using the MAXIscript^TM^ T7 Transcription Kit (Invitrogen). The obtained *nid2a*-exon2-gRNA was extracted using Microspin^TM^ G-50 columns (Sigma). *Cas9* mRNA was generated as reported (86). Injection mixes were prepared with 30ng/µl gRNA and 100ng/µl *Cas9* mRNA in Danieau’s buffer supplemented with 0.05% Phenol Red. Embryos were injected at the one-cell-stage and raised to adulthood. F1 individuals were screened for germline transmission by PCR using the primers 5’-CAGGATCTTCCCATGGAGAA-3’; 5’-CTTTTGACATCCGCTTCTGC-3’ followed by a T7 endonuclease assay (NEB) to detect indels. Subsequent sequencing of indel-harboring amplicons led to the identification of an allele with a 7bp (TTTTTCC) insertion in exon 2 of the *nid2a* gene, leading to a frame shift at amino acid residue (aa) 107 of the normally 1576 aa protein, N-terminal of the first annotated domain (NIDO domain; 113 – 280), and premature truncation of the protein after additional 14 amino acid residues (FFRLPRHRPL SGRH). This allele, which most likely represents a functional null (amorph), was used to establish the stable *nid2a^fr52^* line. Genotyping was performed via PCR of genomic DNA with exon 2 primers, allowing electrophoretic size-discrimination of wild-type and mutant bands on a 3.5% agarose gel in TBE buffer.

Zebrafish and mice were kept under standard conditions, zebrafish embryos were incubated at 28°C in E3 medium (5mM NaCl, 0.17mM KCl, 0.33mM CaCl_2_, 0.33mM MgSO_4_).

Zebrafish and mouse breeding and zebrafish experiments were approved by the national animal care committees (LANUV Nordrhein-Westfalen; 40.15.097; 84-02.04.2012.A251; 84-02.04.2016.A390; 84-02.04.2018.A281; City of Cologne; 576.1.36.6.G13/18 Be) and the University of Cologne (4.16.024; 4.18.013).

### Zebrafish RNA and Morpholino oligonucleotide injections for loss-of-function, rescue and synergistic enhancement studies

Morpholino oligonucleotides (MOs) targeting translational start or splice sites were ordered from Gene Tools (Philomath, OR), dissolved in distilled water to 1 mM stock solutions and kept at room temperature. For injection, aliquots were heated to 65°C, centrifuged and supernatants were diluted in Danieau’s buffer and Phenol Red (0.05%; Sigma) (87). 2.5 nl of MO solution was injected per embryo at the 1-4 cell stage using glass needles pulled on a Sutter needle puller and a Nanoject injection apparatus (Word Precision Instruments). MOs used and their sequences (given 5’-3’) were as follows:

*hmcn1-ATG*: AAA ACG GCG AAG TTA TCA AGT CCA T
*lama5-splice*: AAC GCT TAG TTG GCA CCT TGT TGG C
*nid2a -1.ATG*: TTT TTG GCG AGC AGC TGG TAC ACA T
*nid2a-2.ATG*: CAG CTG GTA CAC ATG CAG CTC GTG A
*nid2a-3.ATG*: AGC ATG CGG GAC TCT GGC TTT GAA T
*nid1b-ATG*: CTG TAA AAC TCC ACA TAC ACG GGC T

Of the three *nid2a* MOs, *nid2a-1.ATG* gave stronger phenotypes than the two others and was used in all shown experiments. To control the efficacy of the *nid2a-1.ATG* MO, anti-Nid2a immunoblotting and immunofluorescence studies were performed. To control its specificity, rescue experiments with co-injected mouse *Nid2* mRNA were performed, which in the targeted 25 bp region contains 15 mismatches compared to the zebrafish *nid2a* cDNA and the morpholino sequence. To generate full-length mouse *Nid2* mRNA, plasmid pUC19-mNidogen2 (63) was cut with *Xba*I, followed by blunting with Klenow and cutting with *Xho*I. The obtained fragment was cloned into the *Stu*I and *Xho*I sites of the pCS2+ expression vector and the resulting plasmid was linearized with *Not*I and transcribed using the SP6 mMessage Machine Kit (Ambion/ThermoFisher). 2.5 nl *mNidogen-*2 mRNA was injected per embryo at concentrations of 200 ng/µl alone or together with 1 ng/µl *nid2a-1.ATG*-MO.

To control the efficacy of the *nid1b-ATG* MO, a 135bp sequence of zebrafish *nid1b* with 81bp of 5’UTR and the first 18 codons of the ORF were PCR amplified from 24hpf cDNA using the primer pair 5’- GGAAGGATCCACTGCAGACCGGGGCTCCTC-3’ (forward) and 5’-GGAAGAATTCCTGGACCGAGACCAGCAGAGC-3’ (reverse). The fragment was cloned via *Bam*HI and *Eco*RI in frame into expression plasmid XLT.GFPLT-CS2+ (gift from Randall Moon; Addgene plasmid # 17098). The integrity and functionality of the construct was verified by sequencing. To obtain capped *nid1b-GFP* fusion mRNA for injections the plasmid was linearized with *Not*I and transcribed using the SP6 mMessage Machine Kit (Ambion/ThermoFisher). 2.5 nl *nid1b-GFP* mRNA was injected at a concentration of 80 ng/µl alone or together with 1 ng/µl *nid1b-ATG*-MO. Injected embryos were assayed for GFP expression at 80% epiboly stage, using a Leica M165FC fluorescence binocular.

### Generation of zebrafish Hmcn1 and Nid2a antibodies

cDNA constructs encoding amino acid residues 1141 – 1576 of zebrafish Nid2a and residues 23-436 of zebrafish Hmcn1 were generated by RT-PCR on total RNA from 24 hpf embryos and cloned with 5′-terminal *Nhe*I and 3′-terminal *Xho*I restriction sites. PCR products were inserted into a modified pCEP-Pu vector containing an N-terminal BM-40 signal peptide and a C-terminal strepII-tag downstream of the restriction sites (82). Recombinant zebrafish Nid2a and Hmcn1 fragments were generated in HEK293-EBNA cells (Invitrogen) and purified as described above for the mouse proteins and used to immunize rabbits or guinea pigs (Pineda Antikörper Service, Berlin, Germany). Obtained antisera were purified by affinity chromatography on a column with purified Nid2a or Hmcn1 fragment coupled to CNBr-activated Sepharose (GE Healthcare). Specific antibodies were eluted with 3M KSCN, and the eluate was dialyzed against PBS, pH 7.4.

### Mouse and zebrafish sectioning and immunofluorescence studies

Mouse skin tissue was fixed for 2 h in 4% paraformaldehyde and then frozen in optimal cutting temperature compound (OCT, Sakura,Torrance,CA) at -80°C. Mouse skeletal muscle tissue (m. gastrocnemius) was immediately embedded in OCT after harvesting, and frozen in liquid nitrogen-cooled isopentane. The frozen samples were cryosectioned at 8 - 12 µm and stored at -20°C. Muscle sections were fixed in 100% EtOH for 15 minutes. After several washes, sections were blocked with 10% sheep serum for 1 hour and then incubated different primary antibodies in blocking solution: guinea pig-anti-mHMCN1 (1653 – 2275 aa) (88), rabbit-anti-mHMCN2 (4429 – 4753 aa) (88), rabbit-anti-mNID2, rabbit-anti-mLaminin-γ1 antiserum (both kindly provided by Takako Sasaki, Oita University, Japan). After overnight incubation at 4°C the sections were again washed with PBS and incubated with the appropriate secondary antibodies conjugated to Alexa 488 or 555 (Molecular Probes, Eugene, OR, USA) at RT for 2 h.

Zebrafish embryos were fixed 2 h in 4% paraformaldehyde, washed extensively in PBS–Triton X-100 (0.5%), blocked in blocking buffer (10% fetal calf serum and 1% DMSO, in PBS–Triton X-100 0.5%), and incubated at 4°C with the primary antibody: rabbit-anti-zfNid2a (this work), guinea pig-anti-zfHmcn1 (this work), rabbit-anti-pan-Laminin (Sigma-Aldrich, L9393), chicken anti-GFP (Invitrogen, A10262). The following secondary antibodies conjugated to Alexa 488, 555 and 647 (Molecular Probes, Eugene, OR, USA) were used in blocking buffer. Confocal images were obtained using a Zeiss LSM 710, 40×/1.1 W Korr LD C-Apochromat, or 20×/0.8 Plan-Apochromat objective and Zen 2.3 SP1 software. Images were processed using Fiji/ImageJ software to obtain maximum intensity projections, and for adjustment of brightness and contrast.

### Stimulated emission depletion (STED) microscopy

Confocal and STED images were acquired using a gSTED microscope (Leica Microsystems) equipped with a white light laser for excitation. A 592 nm depletion laser was used for Alexa 488 and a 775 nm depletion laser for the two other dyes (Abberior STAR 580 and Abberior STAR 635P). A 100x oil immersion objective with a numerical aperture of 1.4 (Leica Microsystems) was used, and the three channels were acquired in a sequential mode. Deconvolution of the images was done using the software Huygens Essential (Scientific Volume Imaging).

### Zebrafish whole mount in situ hybridizations

Embryos were fixed in 4% PFA in PBS overnight at 4°C, and in situ hybridizations were performed as previously described (89) with RNA probes generated from linearized plasmids using the Roche digoxygenin RNA synthesis kit. For *nid1a*, plasmid *pExpress1-zfnid1a* (IMAGp998B0919858Q; NCBI-Acc.-No. EL646124) with a 843 bp *nid1a* cDNA fragment was linearized with *Sma*I and transcribed with T7 RNA polymerase; for *nid1b,* plasmid *pAMP1-zfnid1b* (IMAGp998P0611194Q; NCBI-Acc.-No. EL646124) with a 486 bp *nid1b* cDNA fragment was linearized with *Eco*RI and transcribed with SP6 RNA polymerase, for *nid2a*, plasmid *pBluescript-zfnid2a* (recloned from IRALp962C1766; NCBI-Acc.-No. CK682403) with a 490bp *nid2a* cDNA fragment was linearized with *Sal*I and transcribed with T3 RNA polymerase, for *nid2b*, plasmid *pCRII-zfnid2b* (recloned from IMAGp998A1717616Q; NCBI-Acc.-No. DV594608) with a 778bp *nid2b* cDNA fragment was linearized with *Kpn*I and transcribed with T7 RNA polymerase, for *lama5*, plasmid *pExpress1-zflama5* (IMAGp998B1314694Q, NCBI-Acc.-No. CF348092) with a 736bp *lama5* cDNA fragment was linearized with *Eco*RI and transcribed with T7 RNA polymerase. For *hmcn1*, plasmid *pGEMT-hmcn1* containing a 0.75 kb *hmcn1* cDNA fragment (51) was linearized with *Not*I and transcribed with SP6 RNA polymerase.

### Mouse and zebrafish transmission and immunoelectron microscopy

For morphological examinations via transmission electron microscopy (TEM), zebrafish embryos were fixed in 2% glutaraldehyde and 2% PFA in 0.1M cacodylate buffer (pH 7.4) for 24 h at 4°C. If not immediately processed, they were then stored at 4°C in 0.05% glutaraldehyde in 0.1M cacodylate buffer (pH 7.4). Fixed specimens were washed several times in PBS, post-fixed with 2% osmium tetroxide in 0.1M PBS for 2 h at 4°C, dehydrated in a graded series of ethanol, and transferred in a reverse ethanol series to EPON-araldite (Hundsman Advanced Materials, Derry, NH) (for orientation and embedding. Semithin sections (350 nm) and ultrathin sections (75 nm) were cut with a Reichert-Jung (Leica, Wetzlar, Germany) UCT microtome. Semithin sections for light-microscopy were stained with methylene blue, ultrathin sections were mounted on copper grids coated with formvar (SPI-Chem), contrasted in a Leica EM AX20 contrasting machine with Laurylab Ultrastain 1 (0.5% uranyl acetate, 15 min; Laurylab, Brindas, France) and Laurylab Ultrastain 2 (lead citrate, 8 min), and examined with a Zeiss EM 902 A electron microscope.

Mouse skin was immuno-gold labelled using en bloc diffusion (90) of primary guinea pig anti-mouse Hmcn1 antibodies (88), followed by secondary anti-guinea pig IgG conjugated with 0.8-gold particles. The 0.8-nm gold was subsequently enhanced with additional gold precipitation followed by standard fixation and embedding for transmission electron microscopy.

Zebrafish embryos were stabilized by high pressure freezing (HPF). Briefly, intact embryos were loaded into HPF planchets, with voids surrounding the tissue replaced with 20 % BSA. A second planchet was placed over the first and then the sandwich was immediately frozen in a Leica EMPact 1 High Pressure Freezer (Leica Microsystems). The two halves of the planchet were then separated under liquid nitrogen to expose the tissue within the cavity. Tissues were transferred to the frozen surface of 1% glutaraldehyde with 0.1% uranyl acetate then warmed to -90C. Utilizing a Leica Automated Freeze Substitution device (AFS), temperature was maintained for 1-3 days at -90°C followed by warming at a rate of 1°C/hour to -80°C where the tissues remained for an additional 24 hours prior to warming at a rate of 3°C/hour to -50°C. The tissues were rinsed in 100% acetone at -50°C, warmed at a rate of 1°C per hour to -20°C, then rinsed again in acetone at -20°C for one hour. Tissues were then infiltrated in 1:1, 1:2, then 1:3 acetone:LRWhite embedding media at -20°C, then warmed to 0°C over 3 hours. The temperature within the sample chamber of the AFS was then rapidly raised to +20°C and the tissues transferred to embedding molds. Polymerization in a nitrogen atmosphere was for 24 hours at 60°C. Ultrathin sections cut from LR White embedded zebrafish were mounted on formvar coated nickel slot grids, dried thoroughly, then floated onto 50ul pools of 0.15M Tris-HCl, pH 7.4 (Tris) for 15 minutes, quenched in 0.05M glycine in 0.15M Tris for 60 minutes, blocked in 2% non-fat dry milk with 0.5% ovalbumin and 0.5% fish gelatin for 30 minutes, rinsed in Tris, then incubated in primary antibody diluted 1:10 in Tris for 120 minutes. Grids were then rinsed in Tris for 30 minutes, then in a combination of 5 and 10 nm Goat colloidal gold secondary conjugate diluted 1:10 in Tris for 60 minutes. Finally, grids were rinsed in Tris for 15 minutes, then in distilled water for 15 minutes. Sections were evaluated unstained for background and specific labeling, then contrasted for 4 minutes in 4% Uranyl acetate followed by quarter-strength Reynold’s lead citrate for 15 seconds (91).

### Statistics

Quantitative experiments were repeated at least three times, reaching similar results. Mean values and standard deviations of all individual specimens from one representative or all independent experiments are presented, as specified in the respective figure legends. Statistical analysis was performed using Graph Pad Prism software. For comparison of multiple groups, one-way ANOVA with post-hoc Tukey’s test, for comparison of two groups, an unpaired two-tailed Student’s t-test, was used to determine significance and obtained p-values are mentioned in the respective figure legends.

## Data availability

Data sets not contained within the manuscript can be shared upon request. Contact information: Matthias Hammerschmidt, phone: ++49-221-470 5665

## Acknowledgments

We thank Heike Wessendorf, Petra Comelli, Iris Riedl-Quinkertz and Mojgan Ghilav for excellent technical assistance, and Astrid Schauss and Christian Jüngst from the CECAD Imaging Facility for their help with STED microscopy.

## Author contributions

JZ, TR, SR, DW, SL, HMP, DK, FZ, WB, GS and MH designed experiments. JZ, TR, SR, HMP, DW, SL, DK and WB generated data. JZ, TR, SR, DW, SL, HMP, JH, DK, FZ, WB, GS and MH analyzed and processed data. JZ, GS and MH wrote the manuscript. MH conceptualized and supervised the project.

## Supplementary Data

**Supplementary Figure S1.**
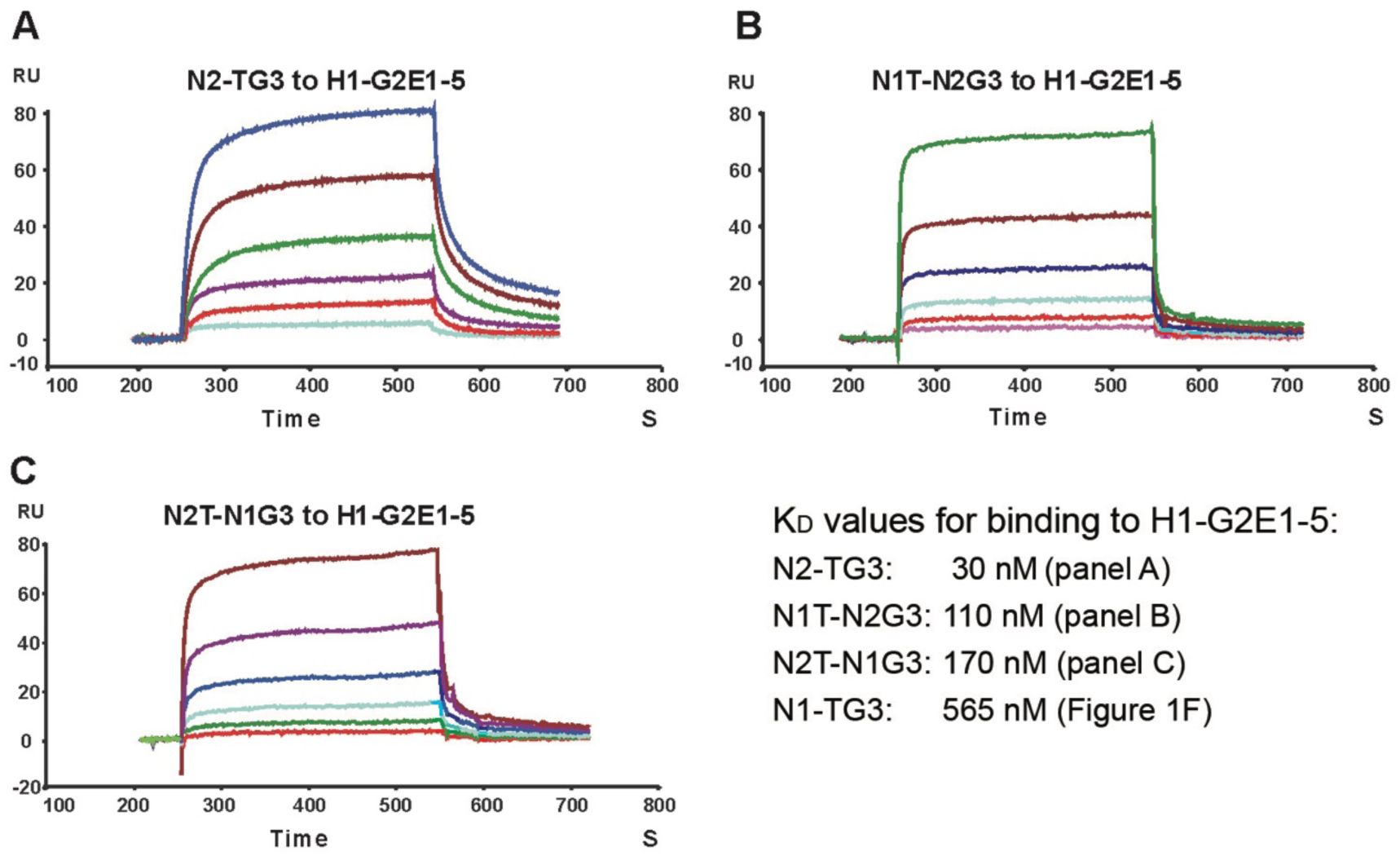
Surface plasmon resonance analysis of the binding between mouse H1-G2E1-5 and hybrid TY-G3 fragments of mouse nidogen-1 and nidogen-2. **A-C:** Sensograms showing the binding of 20 (bottom), 40, 80, 160, 320 and 640 (top) nM Nid2-TG3 (A), 40 (bottom), 80, 160, 320, 640 and 1280 (top) nM N1T-N2G3 (B) or 40 (bottom), 80, 160, 320, 640 and 1280 (top) nM N2T-N1G3 (C) to immobilized H1-G2E1-5 fragment. Abbreviation: RU, resonance unit. K_D_ values calculated from shown sensograms are listed.

**Supplementary Figure S2.**
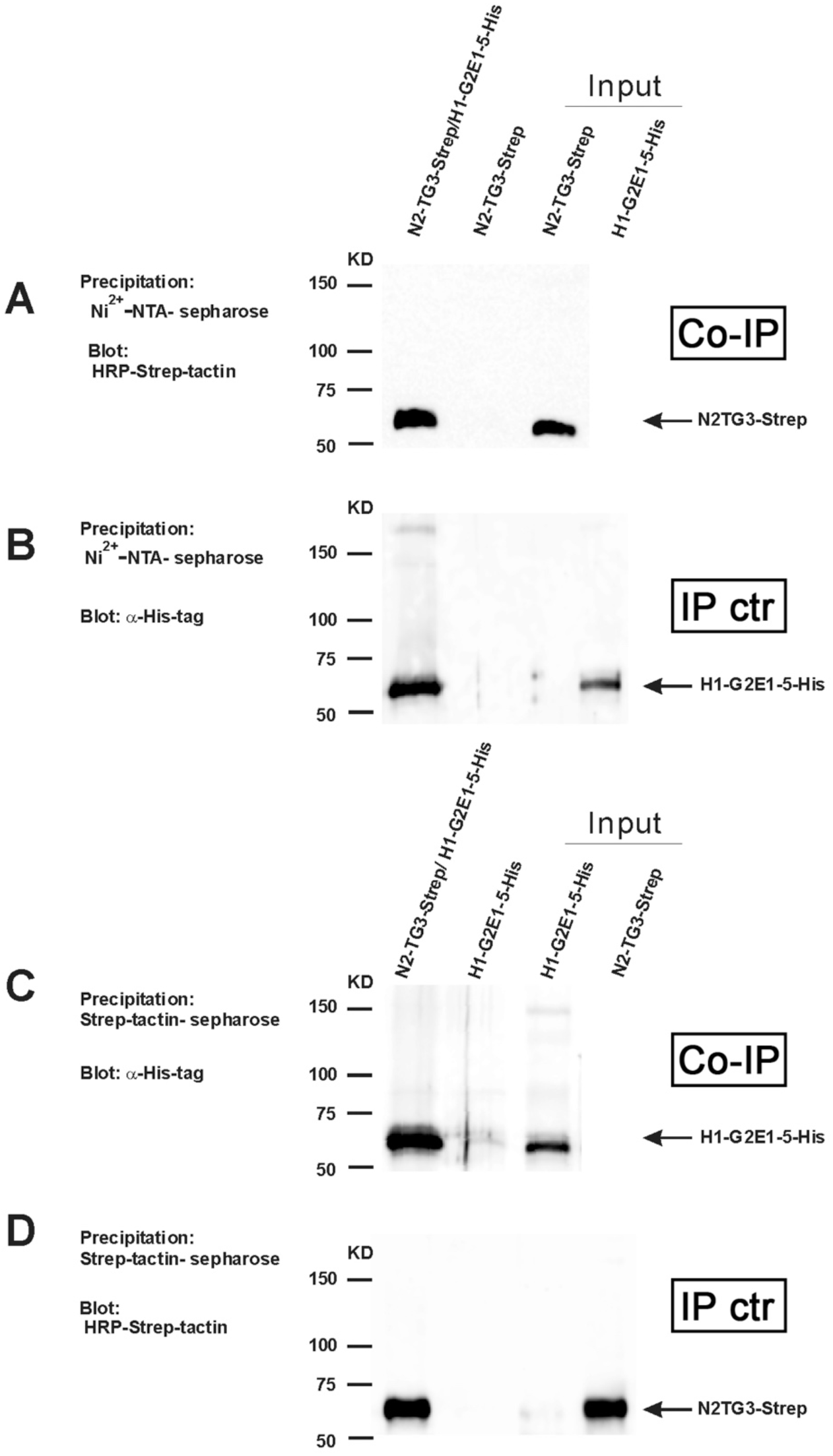
Analysis of the binding between purified mouse NID2 and HMCN1 fragments by co-precipitation. (A,B) Purified N2-TG3-Strep/H1-G2E1-5-His fragments were precipitated by Ni^2+^- NTA sepharose and subjected to western-blotting to detect co-precipitated Strep-tagged protein with HRP-Strep-tactin (A) or, as control of the precipitation, His-tagged protein with anti-His-tag antibodies (B) (lane 1). As negative control, purified N2-TG3-strep alone was precipitated by Ni^2+^-NTA sepharose and detected by western blotting (lane 2). (C,D) For the reverse co-precipitation, purified N2-TG3-Strep/H1-G2E1-5-His was precipitated by Strep-tactin-sepharose and subjected to western-blotting to detect co-precipitated His-tagged protein with anti-His-tag antibodies (C) or, as control of the precipitation, Strep-tagged protein with HRP-Strep-tactin (D) (lanes 1). As negative control, purified H1-G2E1-5-His alone was precipitated by Strep-tactin-sepharose and detected by western blotting (lane 2). Lanes 3-4 of (A-D) are direct loadings of the indicated proteins. Abbreviations: Co-IP, co-immunoprecipitation; IP ctr, control immunoprecipitation.

**Supplementary Figure S3.**
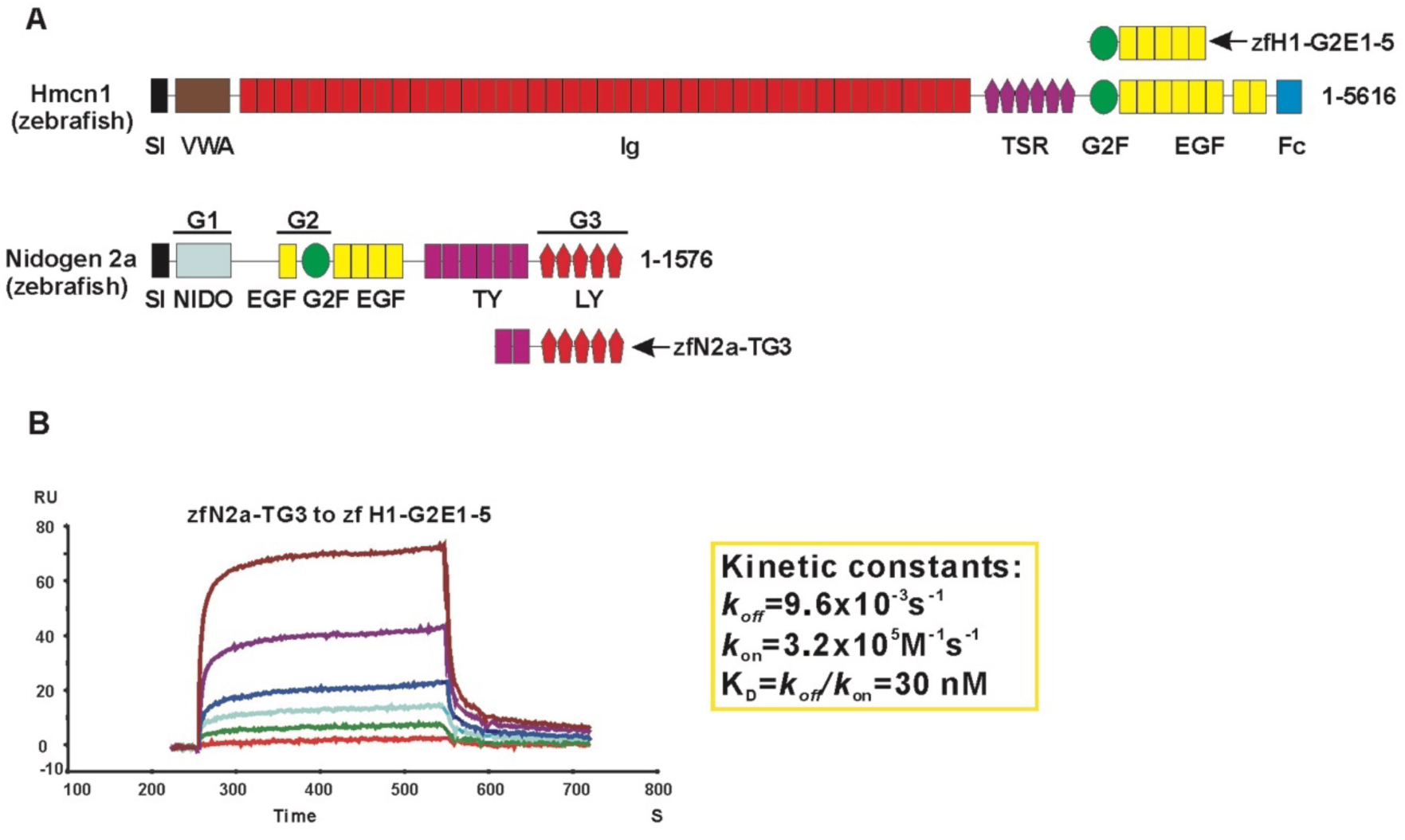
Surface plasmon resonance analysis of the binding between zebrafish H1-G2E1-5 and Nid2a-TG3. **A.** Schematic diagram of zebrafish Hmcn1 and Nid2a proteins and protein fragments used for binding studies. Indicated modules are signal sequence (SI), von Willebrand type A (VWA), immunoglobulin (Ig), thrombospondin (TSR), nidogen G2F (G2F), epidermal growth factor (EGF), fibulin type carboxy-terminal (FCs), Nidogen-like (NIDO), thyroglobulin type-1(TY); and low-density lipoprotein receptor (LY) modules. The globular domains of nidogens are indicated as G1, G2 and G3. Of note, in contrast to mouse NID2 (Figure 1A), zebrafish Nid2a has six, rather than two TY domains N-terminal of its G3 domain. However, according to the UniProt 3D-structures (https://www.uniprot.org/uniprot/X1WE42; https://www.uniprot.org/uniprot/O88322) only the last two of the six TY domains of zebrafish Nid2a are localized between the G2/EGF and G3 domain, similar to the position of the two TY domains of mouse NID2. In contrast, the first four of the six TY domains of zebrafish Nid2a loop out to the opposing side of the protein far remote from the G3 domain and thus are unlikely to contribute to G3-mediated binding. Therefore, we generated a recombinant zfNid2a-TG3 fragment only containing the last two of the six TY domains. B. Sensogram showing the binding of 10 (bottom), 20, 40, 80, 160 and 320 nM (top) zfN2a-TG3 to immobilized zfH1-G2E1-5 fragment. The dissociation constant (*K_D_*) and the kinetic constants *k*_on_ and *k*_off_ used to calculate the *K_D_* value are shown in inset. Abbreviation: RU, resonance unit.

**Supplementary Figure S4.**
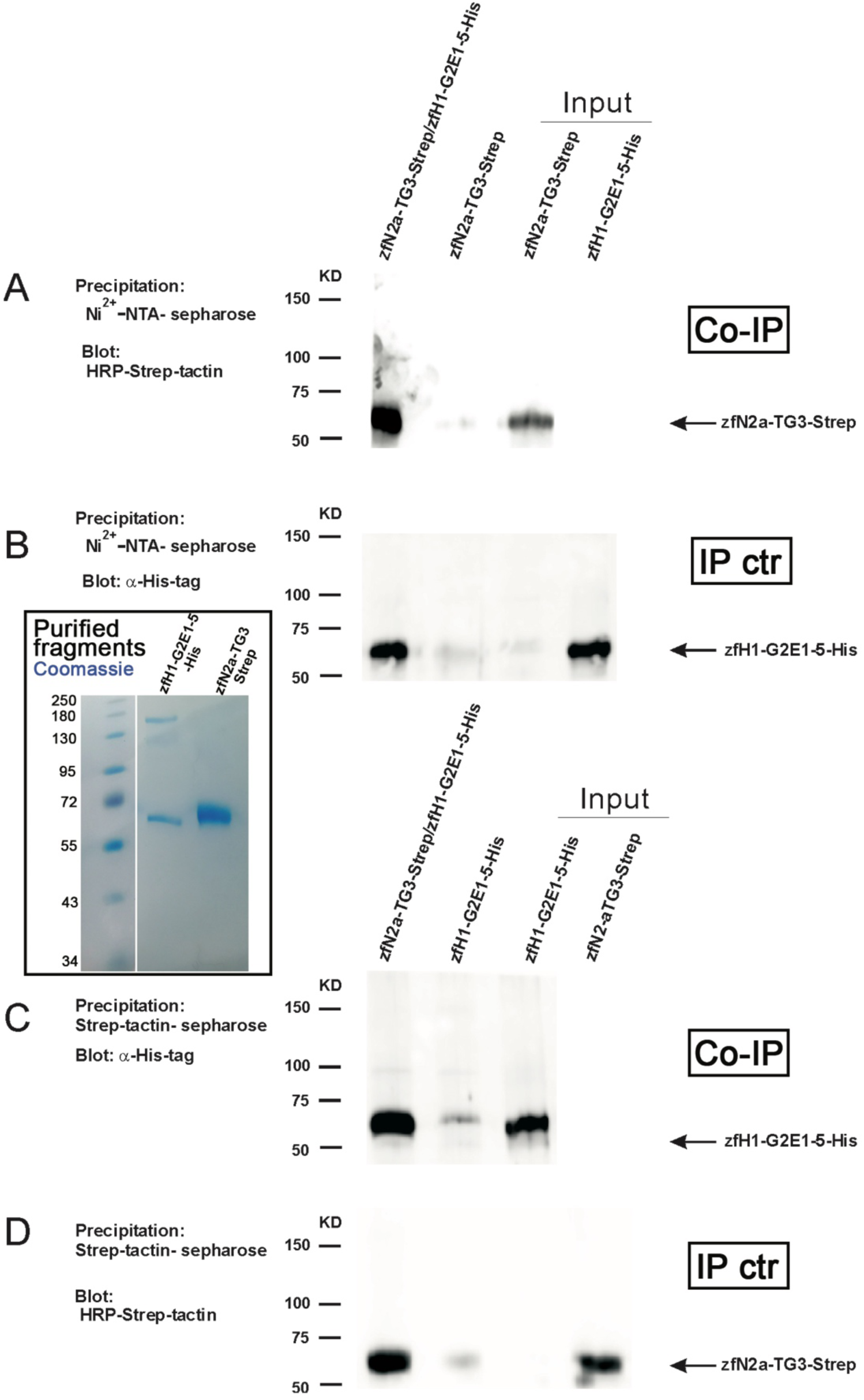
Analysis of the binding between purified zebrafish Nid2 and Hmcn1 fragments by co-precipitation. (A,B) Purified zfN2a-TG3-Strep/zfH1-G2E1-5-His were precipitated by Ni^2+^-NTA sepharose and subjected to western-blotting to detect co-precipitated Strep-tagged protein with HRP-Strep-tactin (A) or, as control of the precipitation, His-tagged protein with anti-His-tag antibodies (B) (lane 1). As negative control, purified zfN2a-TG3-strep alone was precipitated by Ni^2+^-NTA sepharose and detected by western blotting (lanes 2). (C,D) For the reverse co-precipitation, purified zfN2a-TG3-Strep/zfH1-G2E1-5-His was precipitated by Strep-tactin-sepharose and subjected to western-blotting to detect co-precipitated His_6_-tagged protein with anti-His-tag antibodies (C) or, as control of the precipitation, Strep-tagged protein with HRP-Strep-tactin (D) (lanes 1). As negative control, purified zfH1-G2E1-5-His alone was precipitated by Strep-tactin-sepharose and detected by western blotting (lane 2). Lanes 3-4 of (A-D) are direct loadings of the indicated proteins. Inset shows Coomassie blue staining of affinity-purified recombinant zfN2a-TG3-Strep and zfH1-G2E1-5-His fragments. For co-precipitation, relative amounts were adjusted to equimolarity. Abbreviations: Co-IP, co-immunoprecipitation; IP ctr, control immunoprecipitation.

**Supplementary Figure S5.**
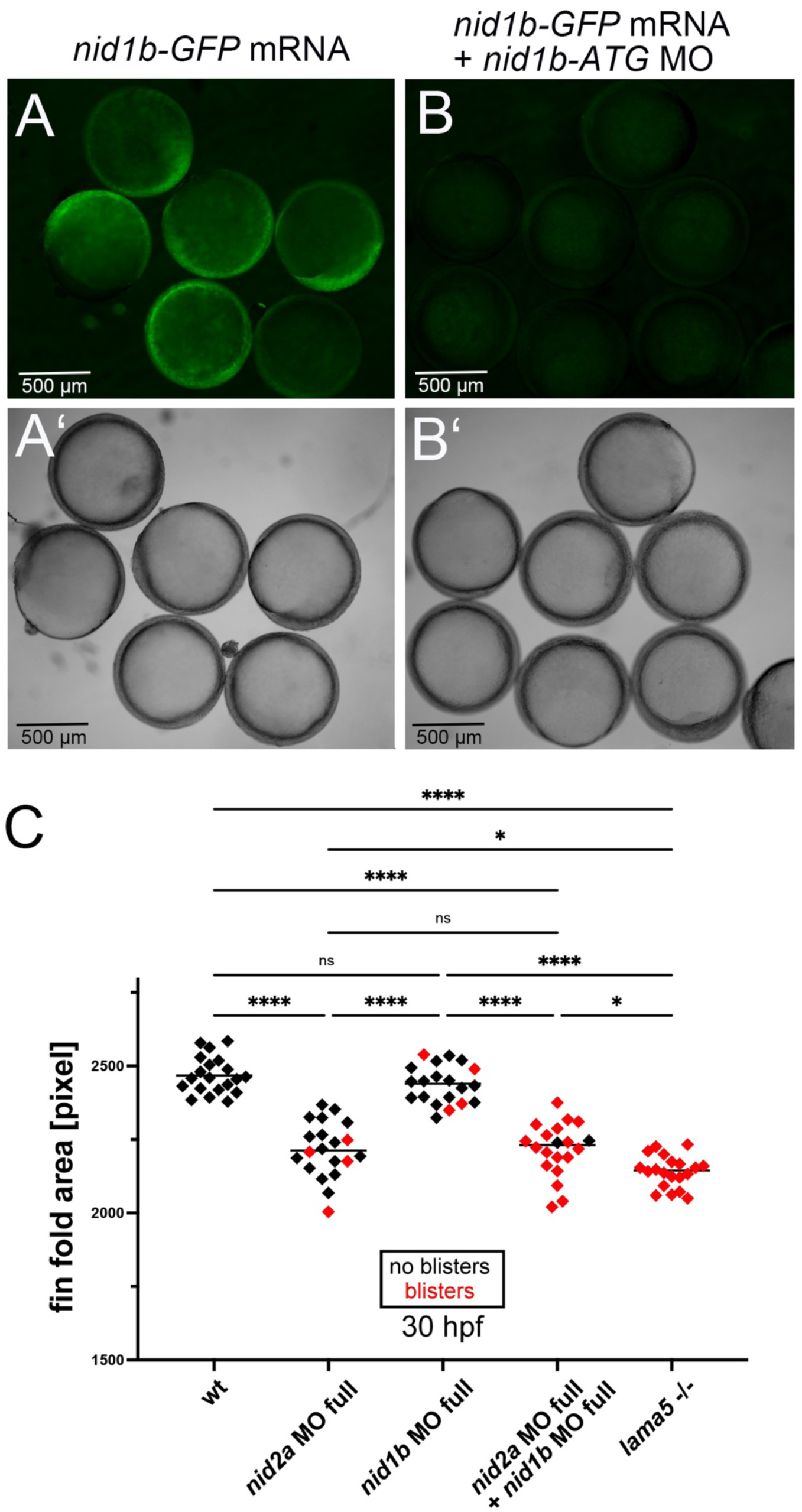
Zebrafish *nid2a* acts in partial functional redundancy with *nid1b*. (A,B) *nid1b-ATG* antisense morpholino oligonucleotide efficiently blocks translation of co-injected mRNA encoding a Nid1a-GFP fusion protein. Panels A,B show green fluorescence (to reveal Nid1a-GFP protein) images of embryos injected with *nid1b-GFP* mRNA alone (A) or co-injected with *nid1b-GFP* mRNA and *nid1b-ATG* MO (B); panels A’,B’ the corresponding bright field images. (C) Scatter plot quantifying median fin fold sizes, indicative of compromised BM-epidermal linkage, in 30 hpf wild-type embryos injected with the indicated MOs (2.2 ng per embryo) and, as positive reference, *lama5* mutants (compare with Figure 6Y). Each diamond represents one individual embryo. Diamonds representing embryos with blisters, indicative of compromised BM-dermal linkage, are in red. In comparison to *nid2a* single morphants, *nid1b* / *nid2a* double morphants display increased rates of fin fold blistering. Significances of differences were determined via ANOVA followed by the Tukey’s post-hoc test. **** indicates a significant difference of p<0.0001, and * of p<0.0178 - 0.0191, respectively. ns indicates no significant differences.

